# MYC shapes ER-mitochondria calcium transfer by directly targeting *ITPR1*: implications for MYC-induced safeguard mechanisms and cancer

**DOI:** 10.1101/2024.08.28.610025

**Authors:** Xingjie Ma, Athanasios Tsalikis, Mathieu Vernier, Kexin Zhu, Jéhanne El Hassani, Dorian Ziegler, Anda Huna, Claire Coquet, Gwladys Revechon, Céline Margand, Flavie Benard, Joanna Czarnecka-Herok, Isabelle Iacono Di Cacito, Catherine Koering, Valéry Attignon, Marjorie Carrere, Florence Jollivet, Bertrand Mollereau, Christophe Vanbelle, Olivier Gandrillon, Benoît Dumont, Céline Delloye-Bourgeois, Hector Hernandez-Vargas, Laura Broutier, Nadine Martin, David Bernard

## Abstract

The MYC and NMYC transcription factors (TFs) play a key role in cell proliferation and are overexpressed in most cancer cells. However, in normal cells their overexpression triggers safeguard mechanisms promoting cell death and cellular senescence, which are bypassed in cancer cells. The mechanisms of action of this TF family are only partially understood. Here, we reveal that in normal cells MYC binds to the Inositol 1,4,5-Trisphosphate Receptor type 1 (*ITPR1)* gene and upregulates its expression, triggering an ER-mitochondria calcium (Ca^2+^) transfer, which is involved in MYC-induced cell death and senescence. Supporting a tumor suppressive role of MYC/ITPR1 axis, *ITPR1* expression is generally decreased in cancer and reactivation of this pathway induces cancer cell death. Nevertheless, some cancer cells, generally expressing high levels of *MYCN* and/or *MYC*, also express high level of *ITPR1*, which correlates with high expression of *BCL2*, encoding an inhibitor of ITPR1. Strikingly, in high-risk *MYCN*-amplified neuroblastoma, *ITPR1* expression is controlled by NMYC and its level correlates with worse patient survival. In these cells, blocking the interaction between BCL2 and ITPR1 induces mitochondrial Ca^2+^ accumulation and cell death, and decreases tumor size. Collectively these data highlight a new function of MYC factors by controlling Ca^2+^ signaling, which could constitute an unsuspected vulnerability for some cancer cells, including high-risk *MYCN*-amplified neuroblastoma cells.

## INTRODUCTION

MYC is a widely expressed transcription factor (TF) that controls the expression of a large panel of target genes regulating multiple pathophysiological processes, including metabolism, proliferation, cell death, cellular senescence, cancer or aging ^1–3^. The MYC TF is required for cell proliferation and exerts its pro-proliferative activity by inducing the transcription of genes encoding positive regulators of cell cycle, such as cyclins, cyclin-dependent kinases and E2F TFs ^4^. Owing to the role of MYC in promoting cell proliferation, its gain-of-function is observed in 70% of cancers, where it acts as a critical driver of cancer formation and progression and can be a therapeutic target ^1,5^. Nevertheless, pathogenic activation of MYC in normal cells initially leads to the activation of safeguard mechanisms, mainly cell death and cellular senescence, that counteract its pro-oncogenic effects. MYC promotes these cytotoxic and cytostatic effects by directly inducing the transcription of pro-apoptotic factors of the BCL2 family, such as BIM, or by inducing the transcription of p14^ARF^, p15^INK4B^ and/or DNA damage, which leads to the activation of p53. Hence, in order to develop, cancer cells must inhibit or circumvent these regulators of apoptosis and senescence ^6,7^. MYC belongs to a family of TFs including NMYC which is primarily expressed in the nervous system during development. MYC and NMYC are considered functionally equivalent, albeit they promote different types of tumors due to their distinct expression profiles ^8–11^. Although several targets and downstream effector pathways are documented, the mechanisms by which MYC TFs induce their effects are only partially understood.

Ca^2+^ levels as well as its dynamics within the cell participate in crucial cellular processes that include cell proliferation, metabolism, cellular senescence and cell death ^12–18^, reminiscent of the broad spectrum of effects induced by MYC ^1,3,19^. Although Ca^2+^ is required for cell viability and functions, cells tightly regulate Ca^2+^ levels and distribution, given that high levels in some organelles are toxic. Indeed, cells maintain low Ca^2+^ concentrations in the cytoplasm, whereas some organelles, such as the endoplasmic reticulum (ER), store it efficiently and release it upon stimulation. ER-mitochondria Ca^2+^ transfer has been described in the process of cell death, during which mitochondrial Ca^2+^ accumulation induces mitochondrial permeabilization and fatal cytochrome c release^17,20^, and was more recently shown to lead to cellular senescence in normal cells ^14,15,18,21–23^. Inositol 1,4,5-Trisphosphate Receptor ITPRs or IP3Rs, a family of 3 members, are channels ensuring release of Ca^2+^ from the ER and thus contributing to the accumulation of Ca^2+^ in the mitochondria, thereby regulating cell death and cellular senescence ^18,23,24^.

Here, we reveal that in normal cells MYC directly binds to the *ITPR1* gene and upregulates its expression, thus promoting ER-mitochondrial Ca^2+^ transfer, cell death and cellular senescence. In cancer cells, the function of ITPR1 is largely lost, either owing to a decrease in its expression, or to an increase in the expression of *BCL2*, encoding an inhibitor of this channel. Strikingly, in *MYCN*-amplified neuroblastomas, with high levels of *ITPR1* and *BCL2* expression, NMYC controls the expression of *ITPR1*. In this context, NMYC/ITPR1/mitochondrial Ca^2+^-dependent cell death can be restored by blocking the inhibitory effect of BCL2 on ITPR1, suggesting a new vulnerability of these high-risk cancers.

## RESULTS

### The MYC transcription factor directly induces *ITPR1* gene expression

Based on the critical role of ITPRs and ER-mitochondria Ca^2+^ transfer in cell death and senescence, we wanted to know whether this signaling, at the crossroad of several cell fates, could be linked to MYC activity. MRC5 normal human fibroblasts were transduced to stably express a 4OHT-inducible MYC ^25^. In this cellular system, MYC activation or knockdown led respectively to an increase or decrease in *ITPR1* mRNA levels, without significantly affecting *ITPR2* and *ITPR3* levels (Figure 1A and Figure S1A). Given the rapid detection of *ITPR1* expression, 6 hours after MYC activation, a similar timeframe to the well-known direct MYC target gene *BIM* ^26^ and much sooner than the induction of the well-known indirect MYC-regulated gene *PUMA* ^27,28^ (Figure 1B), we speculated that *ITPR1* was a direct target for MYC. This was confirmed through ChIP-seq analysis that revealed the binding of MYC on the *ITPR1* promoter in various human cells (Figure 1C) and on the *Itpr1* promoter in mouse cells (Figure S1B). The transcriptional activity of MYC relied on its dimerization with its partner MAX and on the TRRAP co-factor ^29^, as their decrease largely prevented *ITPR1* upregulation by MYC (Figure S1C-D and Table S1). Interestingly, the other MYC co-factors tested, namely KAT2A, KAT5, RUVBL1, RUVBL2 and SUPT5H, did not contribute to the upregulation of *ITPR1* by MYC (Table S1). Together these results support that *ITPR1* is a direct target gene of MYC, which induces its expression.

**Figure 1.**
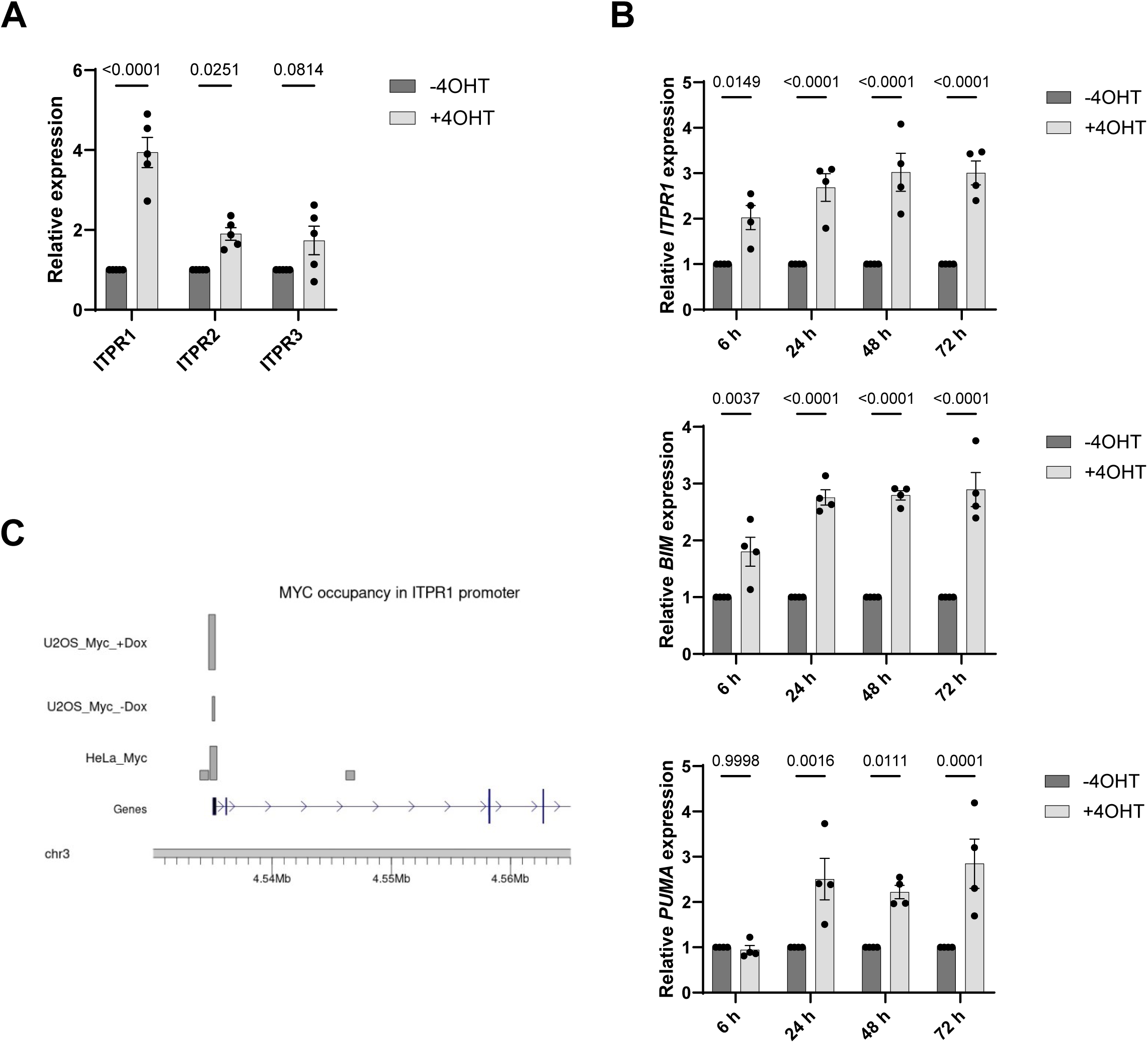
*ITPR1* expression is directly induced by MYC. **A.** RT-qPCR of *ITPR1*, *ITPR2* and *ITPR3* genes at day 3 after treatment of MRC5-MYC:ER cells with 4OHT. Mean +/- SEM of n = 5 independent experiments. Two-Way ANOVA. P-values are indicated. **B.** RT-qPCR of *ITPR1*, *BIM* and *PUMA* genes at the indicated times after treatment of MRC5-MYC:ER cells with 4OHT. Mean +/- SEM of n = 3 independent experiments. Two-Way ANOVA. P-values are indicated. **C.** ChIP-seq analysis of MYC occupancy at the *ITPR1* promoter region in MYC-overexpressing cells (U2OS, HeLa) (GSE44672).

### ITPR1 mediates MYC-induced safeguard mechanisms

We then wondered whether ITPR1 contributed to MYC-induced cell death and/or senescence in normal cells. We first validated that, in addition to *ITPR1* mRNA, the level of the ITPR1 protein was also regulated by MYC (Figure 2A-B). As expected, MYC activation led to a drop in the number of cells, as seen by crystal violet staining, and we observed that this drop was largely prevented by the knockdown of ITPR1 (Figure 2C), but not of ITPR2 or ITPR3 (Figure S2A). To better understand the underlying mechanisms, we then quantified the number of dying cells, through a trypan blue assay or live imaging using SYTOX dye, and the number of senescent cells using the senescence-associated-β-galactosidase (SA-β-Gal) activity assay. Knocking down ITPR1 upon MYC activation impeded both MYC-induced cell death (Figure 2D-E) and MYC-induced senescence (Figure S2B). Hence, these results indicate that direct upregulation of *ITPR1* expression by MYC contributes to the activation of safeguard mechanisms in normal cells.

**Figure 2.**
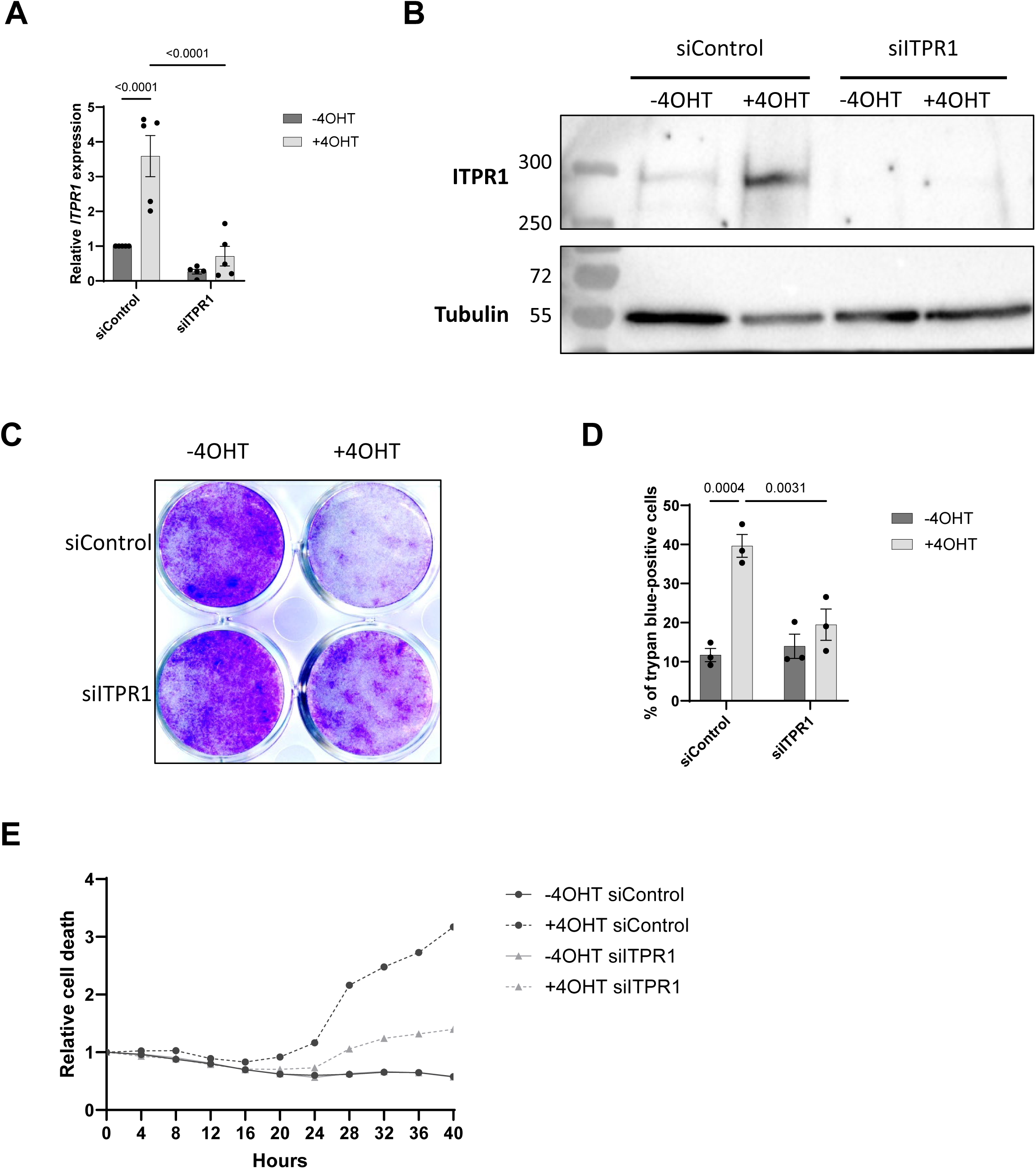
ITPR1 contributes to MYC-induced safeguard mechanisms. MRC5-MYC:ER cells were transfected with a control siRNA pool (siControl) or with a siRNA pool targeting ITPR1 (siITPR1) and then treated or not with 4OHT. **A.** RT-qPCR of *ITPR1* gene at day 3 after 4OHT treatment. Mean +/- SEM of n = 5 independent experiments. Two-Way ANOVA. P-values are indicated. **B.** Western blot showing ITPR1 protein levels at day 3 after 4OHT treatment and α-Tubulin protein levels as loading control. Representative images of n = 3 independent experiments. **C**. Crystal violet staining at day 10 after 4OHT treatment. Representative image of n = 3 independent experiments. **D**. Percentage of trypan blue-positive cells counted at day 3 after 4OHT treatment. Mean +/- SEM of n = 3 independent experiments. Two-Way ANOVA. P-values are indicated. **E**. Cell death monitored using SYTOX Green in the first 40h after 4OHT treatment. Representative experiment from n=3 independent experiments.

### ITPR1-dependent ER-mitochondria Ca^2+^ transfer is involved in MYC-induced safeguard mechanisms

Given that Ca^2+^ release through ITPRs channels and its subsequent accumulation in the mitochondria can contribute to cell death and cellular senescence ^18,23,24^, we assessed mitochondrial Ca^2+^ accumulation in MYC-activated cells using a mitochondria-targeted ratiometric Ca^2+^ probe ^21,23^. Strikingly, MYC induced a rise in mitochondrial Ca^2+^, which was inhibited by knocking down ITPR1 or VDAC3, a channel involved in Ca^2+^ entry into the mitochondria (Figure 3A-B). This increase in mitochondrial Ca^2+^ induced by the MYC/ITPR1 axis was critical for MYC-induced cell death and cellular senescence, as both cell fates were inhibited after VDAC3 knockdown, similarly to ITPR1 knockdown, during MYC activation (Figure 3C and S3). Altogether, in response to MYC activation, ITPR1 and mitochondrial Ca^2+^ accumulation mediate cell death and cellular senescence, two cell fates preventing tumorigenesis.

**Figure 3.**
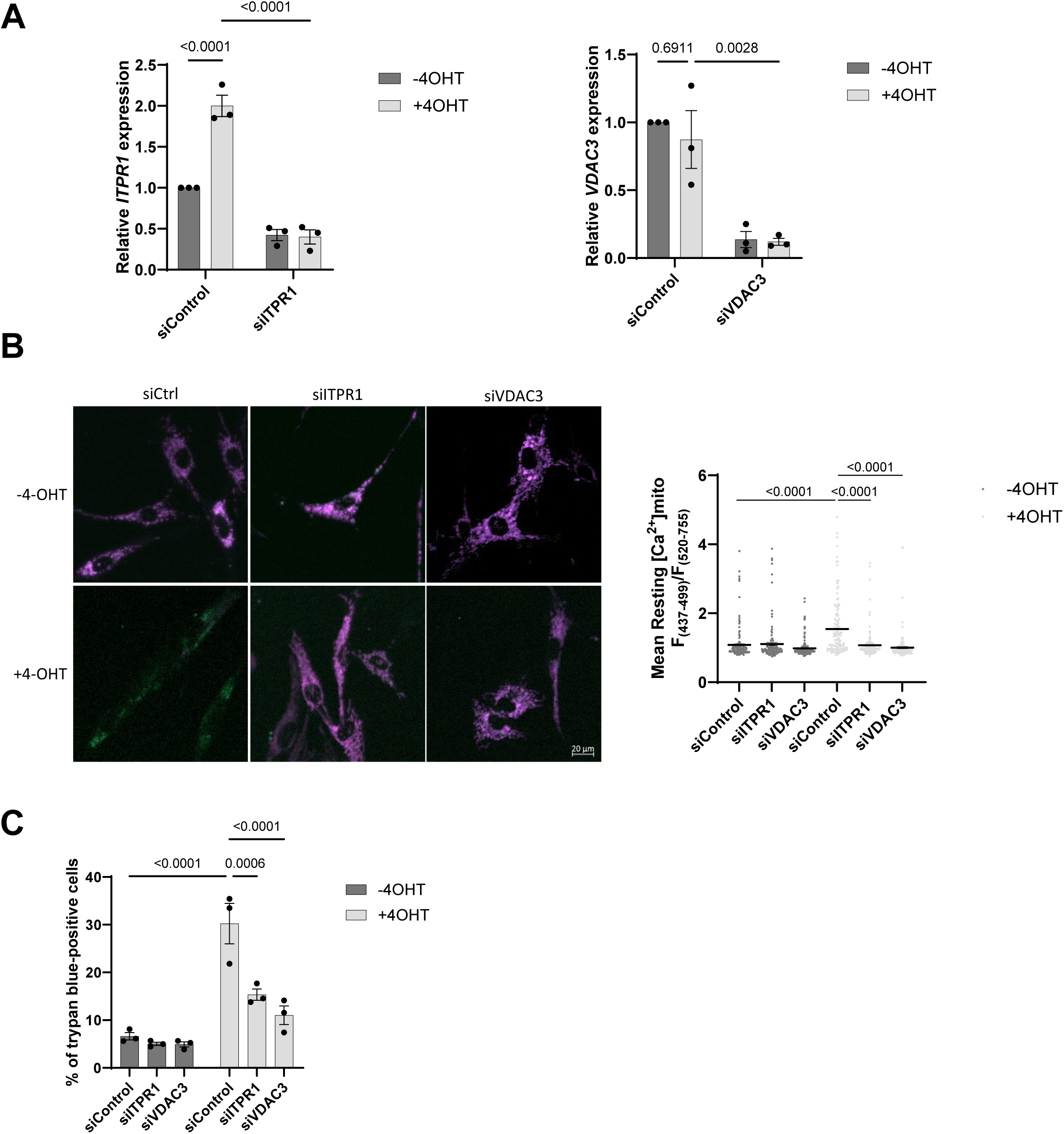
ITPR1-dependent mitochondrial calcium accumulation contributes to MYC-induced safeguard mechanisms. **A.** MRC5-MYC:ER cells were transfected with a control siRNA pool (siControl) or with a siRNA pool targeting ITPR1 (siITPR1) or VDAC3 (siVDAC3) and then treated or not with 4OHT. RT-qPCR of *ITPR1* gene (left) or *VDAC3* gene (right) at day 3 after 4OHT treatment. Mean +/- SEM of n = 3 independent experiments. Two-Way ANOVA. P-values are indicated. **B**. MRC5-MYC:ER cells overexpressing mito-GEM-GECO1 ratiometric mitochondrial Ca^2+^ gene reporter were transfected with a control siRNA pool (siControl) or with a siRNA pool targeting ITPR1 (siITPR1) or VDAC3 (siVDAC3) and then treated or not with 4OHT for 2 days. Resting mitochondrial Ca^2+^ levels according to the ratio F(437-499)/F(520-755) were measured. Representative images are shown as well as quantitative results from n = 3 independent experiments. siControl -4OHT: n = 148 cells, siITPR1 -4OHT: n = 129 cells, siVDAC3 - 4OHT: n = 198 cells, siControl +4OHT: n = 122 cells, siITPR1 +4OHT: n = 152 cells, siVDAC3 +4OHT: n = 159 cells. Mean ± SEM are shown. Two-way ANOVA test. P-values are indicated. **C**. Percentage of trypan blue-positive cells counted in MRC5-MYC:ER cells transfected with siControl, siITPR1 or siVDAC3 and then treated or not with 4OHT for 3 days. Mean +/- SEM of n = 3 independent experiments. Two-Way ANOVA. P-values are indicated.

### *ITPR1* is largely underexpressed in cancers, and restoring the MYC/ITPR1 tumor suppressive pathway kills cancer cells

A gain-of-function of MYC is observed in most of cancers, stemming directly from gene mutations and amplifications, or indirectly through the activation of upstream oncogenic pathways and partners ^5^. As our results support a critical, anti-tumoral role for ITPR1 by promoting cell death and cellular senescence in response to MYC activation, we hypothesized that ITPR1 may be lost in many tumors. We examined its expression profile in human cancers using TCGA datasets and observed that *ITPR1* expression was significantly lower in most types of cancers compared to their normal counterparts (Figure 4A). We then forced the activation of the MYC/ITPR1 pathway in U2OS cancer cells, in which *ITPR1* expression levels are low according to the CCLE database. This led to a decrease in the number of cells and an increase in cell death, effects that were reverted by knocking down ITPR1, further supporting its tumor suppressive role (Figure 4B-E). Collectively, these data show that *ITPR1* expression is low in most human cancers and that forcing the MYC/ITPR1 axis induces cancer cell death.

**Figure 4.**
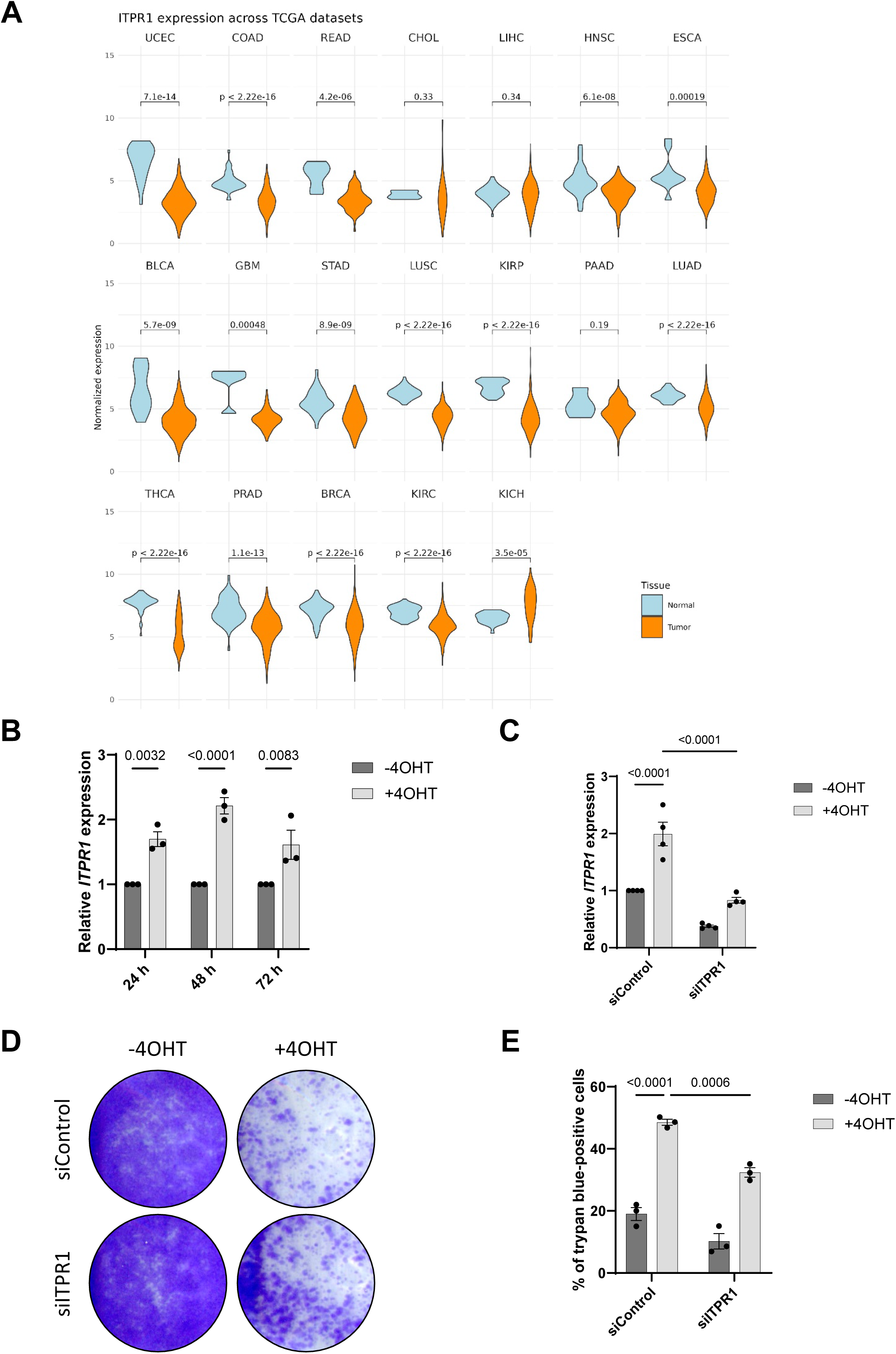
*ITPR1* expression is frequently decreased in cancer and restoring MYC/ITPR1 axis in cancer cells exerts cytotoxic activity. **A.** Violin plots depicting the distribution of *ITPR1* expression levels in normal tissue samples compared to tumor tissue samples across various cancer types from The Cancer Genome Atlas (TCGA) dataset. Wilcox test. P-values are indicated. **B-E**. U2OS cancer cells were infected with a MYC:ER encoding retroviral vector and puromycin selected. **B.** RT-qPCR of *ITPR1* gene at the indicated times after 4OHT treatment. Mean +/- SEM of n = 3 independent experiments. Two-Way ANOVA. P-values are indicated. **C-E**. U2OS-MYC:ER cells were transfected with a control siRNA pool (siControl) or with a siRNA pool targeting ITPR1 (siITPR1) and then treated or not with 4OHT. **C**. RT-qPCR of *ITPR1* gene 3 days after 4OHT treatment. Mean +/- SEM of n = 4 independent experiments. Two-Way ANOVA. P-values are indicated. **D**. Cell density assessed by crystal violet staining 6 days after 4OHT treatment. Representative of n=3 experiments. **E**. Percentage of trypan blue-positive cells 6 days after 4OHT treatment. Mean +/- SEM of n = 3 independent experiments. Two-Way ANOVA. P-values are indicated.

### *ITPR1* expression is high in some cancer cells with high *BCL2* and high *MYC* or *MYCN* expression levels, especially in *MYCN*-amplified neuroblastomas

Even though *ITPR1* expression was relatively low in tumor tissues compared to normal tissues (Figure 4A), its range of expression in cancer cells was broad (Figure 5A). Given that BCL2 and BCL2L1 (BCL-xL) anti-apoptotic factors are known inhibitors of the ITPR1 channel ^30,31^, we wondered whether cancer cells expressing high levels of *ITPR1* also displayed high levels of *BCL2* and/or *BCL2L1* expression. *BCL2* expression, but not *BCL2L1* expression, was positively correlated with *ITPR1* expression (Figure 5A). This suggests that cancer cells displaying high level of *ITPR1* expression may have developed a strategy to decrease ITPR1 activity by upregulating BCL2. We then selected the cancer cell lines with the highest *BCL2* (>3) and *ITPR1* (>3) expression levels (Figure 5A and Table S2). These 54 cancer cell lines were predominantly derived from autonomic ganglia generating neuroblastoma, and from hematopoietic and lymphoid tissues, representing a wide spectrum of different types of cancers (Figure 5B), and were enriched in cancer cell lines displaying either high levels of *MYCN or MYC* expression or both (Figure 5C). For further experiments we focused on neuroblastoma cells lines that were all displaying *MYCN* amplification. Interestingly, in human neuroblastoma samples, *ITPR1* and *BCL2* expression levels were strongly or weakly correlated in *MYCN*-amplified (Figure 5D) or non-amplified neuroblastoma, respectively (Figure S4). Taken together, these observations suggest that *ITPR1* expression can be sustained in some cancer cells, mostly in the context of high *MYCN* and/or *MYC* expression (especially in the case of *MYCN*-amplified neuroblastomas), as well as of high *BCL2* expression.

**Figure 5.**
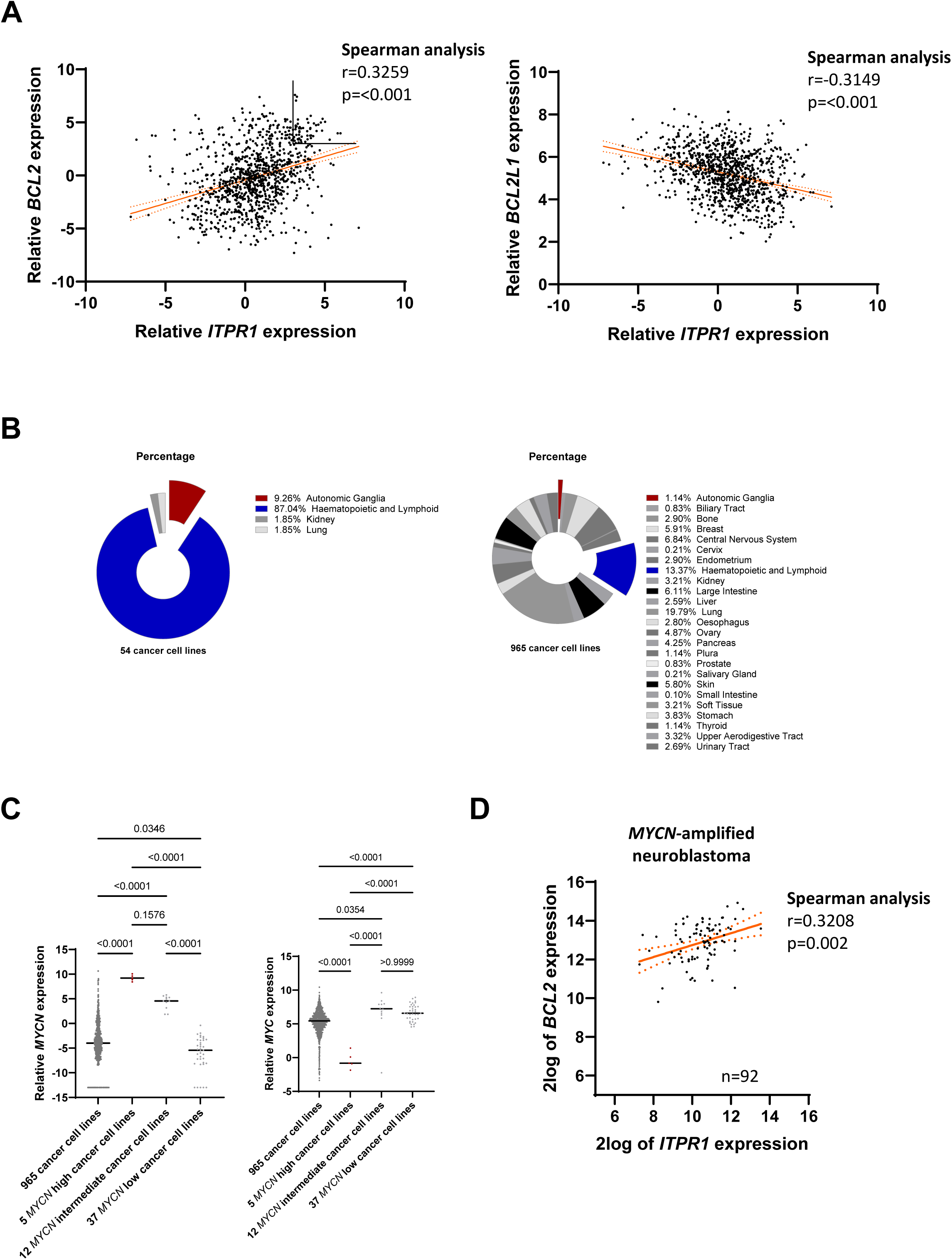
*ITPR1* expression is high in some tumors with high *BCL2* expression, especially in *MYCN*-amplified neuroblastomas. A-C. Expression of *ITPR1*, *BCL2* and *BCL2L1* (BCL-xL) genes in 1,019 cancer cell lines extracted from the Cancer Cell Line Encyclopedia. **A**. Correlation between *ITPR1* and *BCL2* or *BCL1L1* levels of expression. The cancer cell lines displaying the highest expression levels of *ITPR1* (>3) and of *BCL2* (>3) were selected from the left-hand graph (upper right square: 54 cancer cell lines). **B.** A circular diagram showing the distribution of the cancer cell lines according to their tissue of origin in the 54 cancer cell lines displaying high *ITPR1* and *BCL2* expression levels (left) and in the other 965 cancer cell lines (right). **C.** Expression level of *MYCN* and *MYC* in these cancer cell lines. The 54 cancer cell lines were split into 2 categories: *MYCN* high, and *MYCN* intermediate or *MYCN* low. One-Way ANOVA. **D.** Expression data of *BCL2* and *ITPR1* extracted from a human neuroblastoma dataset (Tumor Neuroblastoma – SEQC – 498 – custom – ad44kcwolf) using the R2 Genomics Analysis and Visualization Platform. Correlation of expression between *BCL2* and *ITPR1* mRNA levels is shown in 92 *MYCN*-amplified neuroblastomas.

### NMYC binds to *ITPR1* and activates its expression in *MYCN*-amplified neuroblastomas

Next, we wondered whether NMYC could control the expression of *ITPR1* in *MYCN*-amplified neuroblastomas. We found that NMYC can bind the same region in Kelly neuroblastoma-derived cells than MYC in various human cells (Figure 6A). Chemical inhibition of NMYC using VPC-70619 ^32^ or shRNA significantly decreased *ITPR1* expression (Figure 6B and S5A). We then confirmed these results using another *MYCN*-amplified neuroblastoma-derived cell line, SHEP, in which *MYCN* expression could be shut down using a TET-OFF system ^33,34^. The shutdown led to a decrease in (i) *MYCN* mRNA levels (Figure S5B), (ii) the binding of NMYC on the *ITPR1* promoter (Figure 6C), and (iii) the level of *ITPR1* mRNA (Figure 6D). Finally, *MYCN* and *ITPR1* mRNA levels were strongly correlated in *MYCN*-amplified human neuroblastoma samples and not in *MYCN* non-amplified neuroblastomas (Figure 6E). Hence, these results support that NMYC directly controls the expression of *ITPR1* in *MYCN*-amplified neuroblastomas.

**Figure 6.**
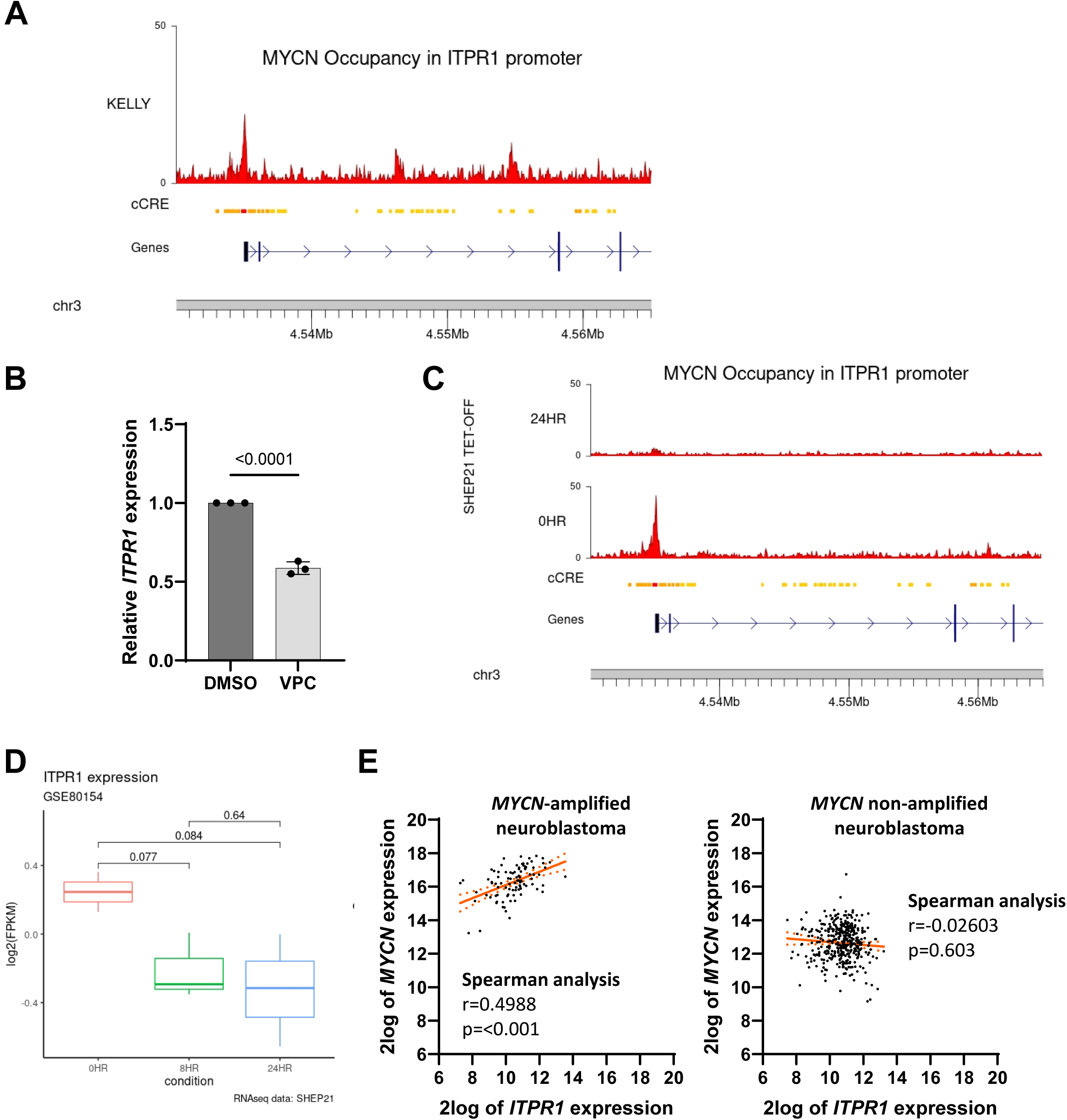
NMYC binds *ITPR1* and activates its expression in *MYCN*-amplified neuroblastoma cells. **A.** Representation of NMYC ChIP-seq peaks on the *ITPR1* promoter in the Kelly *MYCN*-amplified neuroblastoma cell line (GSE80151). The cCRE track summarizes the ENCODE Candidate Cis-Regulatory Elements (cCREs) combined from all cell types with red indicating promoter and orange and yellow indicating proximal and distal enhancer-like signatures, respectively. **B.** RT-qPCR of *ITPR1* gene in Kelly cells treated with DMSO (negative control) or NMYC inhibitor VPC-70619. Mean +/- SEM of n = 3 independent experiments. T-test. P-values are indicated. **C.** ChIP-seq analysis of NMYC occupancy at the *ITPR1* promoter region in neuroblastoma cells expressing or not NMYC (SHEP21 TET-OFF system). **D.** RNA-seq analysis of *ITPR1* expression level after 24 h of NMYC depletion in SHEP21 TET-OFF system. Wilcoxon test. P-value is shown. **E.** Expression data of *MYCN* and *ITPR1* extracted from a human neuroblastoma dataset (Tumor Neuroblastoma – SEQC – 498 – custom – ad44kcwolf) using the R2 Genomics Analysis and Visualization Platform. Correlation of expression between *MYCN* and *ITPR1* is shown in 92 *MYCN*-amplified neuroblastomas (left panel) and in 401 *MYCN* non-amplified neuroblastomas (right panel).

### *MYCN*-amplified neuroblastomas with the strongest *ITPR1* expression are of worse prognosis but are vulnerable to ITPR1 activation by inhibiting ITPR1/BCL2 interaction

Given the role of the NMYC/ITPR1 pathway in neuroblastomas, we wondered whether it was correlated with the survival of patients with either *MYCN*-amplified or *MYCN*-non-amplified neuroblastomas. As expected, patients with *MYCN*-amplified neuroblastomas displayed a shorter survival time (Figure 7A and S6A) ^35^. Strikingly high *ITPR1* expression was correlated with a worse survival in *MYCN*-amplified neuroblastomas (Figure 7A), but with a better survival in *MYCN*-non-amplified neuroblastomas (Figure S6A). As higher levels of *ITPR1* expression in *MYCN*-amplified neuroblastomas correlated with worse survival and *ITPR1* expression levels correlated with *BCL2* expression levels, we reasoned that *MYCN*-amplified neuroblastoma cells might be highly sensitive to the release of ITPR1 inhibition by BCL2. To test this hypothesis, several *MYCN*- and *MYCN*-non-amplified neuroblastoma cells were treated with Bird2 peptide, a sequence corresponding to the ITPR1 domain disrupting the ITPR1/BCL2 interaction ^31^, or a control peptide (Table 1). All the 4 *MYCN*-amplified neuroblastoma cell lines were sensitive to Bird2 treatment whereas only 1 out of the 4 *MYCN*-non-amplified neuroblastoma lines was sensitive (Table 1). These results were further confirmed as *MYCN*-amplified Kelly neuroblastoma cells died rapidly after adding Bird2 whereas *MYCN*-non-amplified SKNAS neuroblastoma cells were resistant to this treatment (Figure 7B-C and S6B). As expected, the death of Kelly cells induced by Bird2 was correlated with an increase in mitochondrial Ca^2+^ (Figure 7D). To further confirm this result *in vivo*, we grafted Kelly cells stably expressing the GFP into HH14 chick embryos, in the dorsal roof of the neural tube between somites 18 and 24 to model the formation of neuroblastoma tumors in developing sympathetic tissues as previously reported ^36^. We then injected the control or Bird2 peptide intravenously, and analyzed engrafted embryos at HH25 by 3D imaging (Figure 7E), as previously reported ^37^. Bird2 treatment strongly decreased the size of neuroblastoma tumors (Figure 7F). These data thus reveal that high *ITPR1* expression levels are correlated with a worse prognosis in high-risk *MYCN*-amplified neuroblastomas, and that these neuroblastomas may be sensitive to the inhibition of the BCL2/ITPR1 interaction.

**Figure 7.**
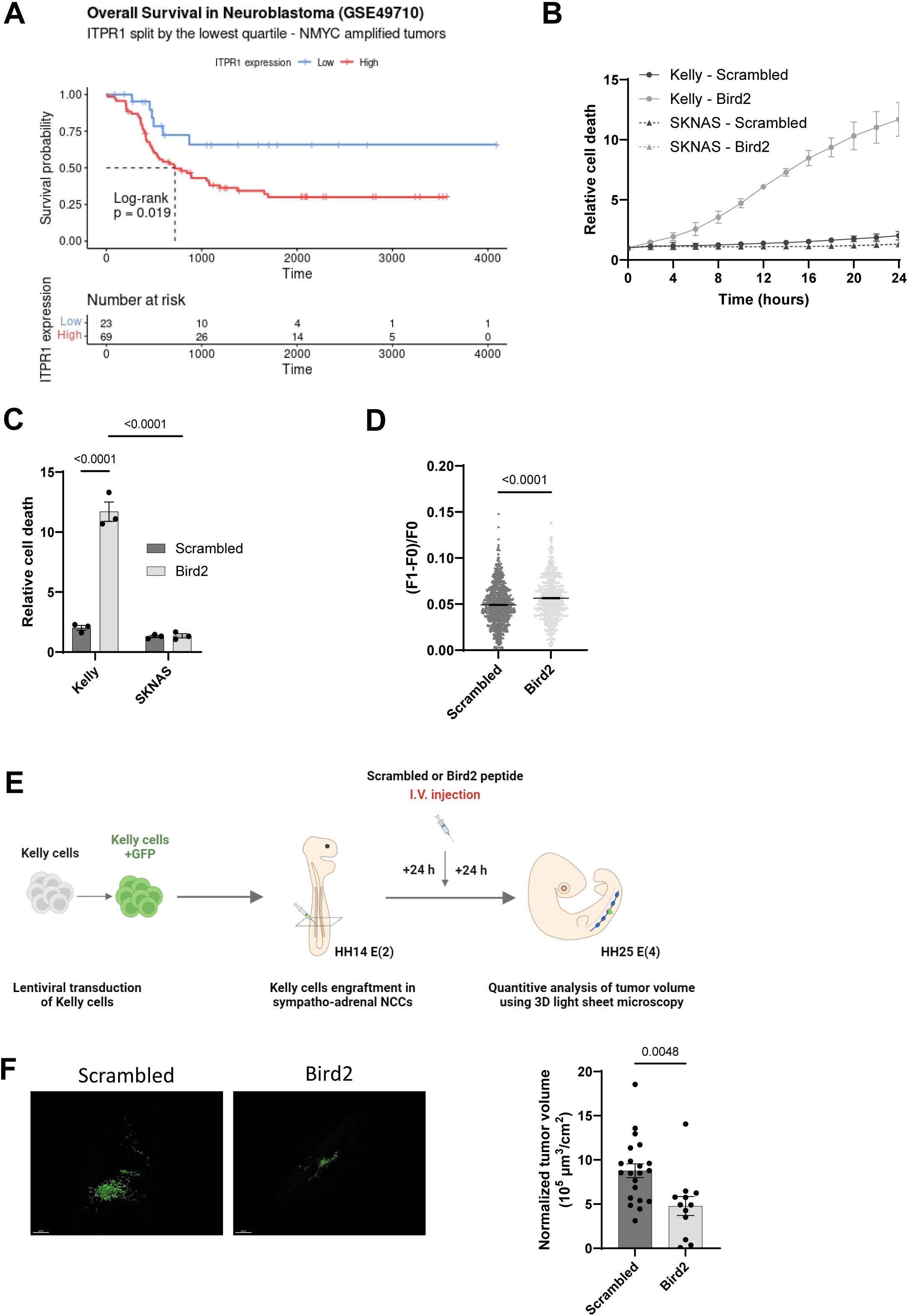
High expression levels of *ITPR1* are of bad prognosis in *MYCN*-amplified neuroblastoma and targeting ITPR1-BCL2 interaction can kill *MYCN*-amplified neuroblastoma cells. **A.** Kaplan-Meier survival curves drawn from GSE49710 dataset from patients with *MYCN*-amplified neuroblastoma. The 2 groups were formed according to the lowest quartile of *ITPR1* expression. **B-C**. Cell death in Kelly and SKNAS neuroblastoma cells monitored using SYTOX Green in the first 24 h after treatment with Scrambled or Bird2 peptides. **B**. Representative curve over a 24 h course. Mean +/- SD of n=3 experiments. **C.** Analysis after 24 h treatment with the peptides. Mean +/- SEM of n = 3 independent experiments. Two-Way ANOVA. P-values are indicated. **D**. Quantification of mitochondrial calcium levels in Kelly cells overexpressing Mitycam, a mitochondrial calcium genetic sensor, in response to treatment with Scrambled or Bird2 peptides. The ratio (F1-F0)/F0 (F1: measurement 3 seconds after injection and F0: measurement 3 seconds before injection) was calculated. n = 3 independent experiments. Scrambled: n = 1,480 cells, Bird2: n = 913 cells. Mean ± SEM are shown. T-test. P-values are indicated. **E.** Schematic representation of the protocol used to assess Bird2 efficacy on neuroblastoma using a model of graft in the chick embryo. **F.** Size of the tumor in the embryo measured through GFP fluorescence. A representative image of neuroblastoma for each condition is shown (left) and quantification of the normalized neuroblastoma volume is displayed for each embryo (right). Scrambled peptide n=21, Bird2 peptide n=12. Mean +/- SEM. T-test. P-values are indicated.

**Table 1.** Sensitivity of different neuroblastoma-derived cell lines to Bird2 peptide according to their MYCN amplification status. The sensitivity to Bird2 peptide when compared to control Scrambled peptide was examined 48 hr after adding 10 mM of the peptides.

| Neuroblastoma-derived cell lines |  | Sensitivity to Bird2 treatment |
| --- | --- | --- |
| MYCN-amplified neuroblastomas | Kelly | YES |
|  | KPNYN | YES |
|  | CLB BERLud 1 | YES |
|  | CLB BAR Rec | YES |
| non MYCN-amplified neuroblastomas | SHSY5Y | NO |
|  | SKNAS | NO |
|  | SKNSH | NO |
|  | CLB-Ga | YES |

To assess the efficiency of the Bird2 therapeutic strategy in a pre-clinical model that can accurately predict outcomes in the clinic ^38^, we established neuroblastoma organoid cultures. Based on previously described neuroblastoma tumor-initiating cell lines and organoids derivation protocols ^39,40^, we established 3D neuroblastoma organoids (NB_O) from *MYCN*-amplified patient-derived xenograft (NB1 and NB2) and human biopsy (NB3) fresh tissues (Figure 8A) as early as 3 days post culture-initiation. NB_O were split every 10 days (600-800µm), at a 1:2 to 1:4 ratio, for at least 6 months. These 3D models were biobanked and reanimated for further cellular and molecular analyses, with a success rate of 100% (n = 3/3). One of the essential points to meet the definition of tumor-derived organoid is the preservation of the cytoarchitecture of the original tissue. Thus, we further validated our models at histological level after 2 to 3 months in culture (Figure 8B). This phenotypic analysis revealed that NB_O preserved the histological features of their primary tumor (Figure 8B), including the expression of sympatho-adrenal markers observed in their parental tumors, as assessed by PHOX2B and CD56 stainings (Figure 8B). The robustness of patient-derived organoids also depends on their ability to preserve the molecular characteristics of their original tumor. We then performed transcriptomic characterization to assess the molecular proximity of NB_O to their corresponding tumor tissue. PCA and Pearson’s correlation heatmaps established from transcriptomic profiling unveiled that NB_O were grouped with their respective tumor-of-origin even after long-term culture and cryopreservation/revival (Figure 8C-D). Hierarchical clustering analysis based on neuroblastoma sympatho-adrenal lineage-related markers ^41^ confirmed the high level of similarities between NB_O and their corresponding tumor samples, even after cryopreservation, while unveiling notable differences between the NB_O lines reflecting the overall neuroblastoma inter-tumoral heterogeneity (Figure 8E-F).

**Figure 8.**
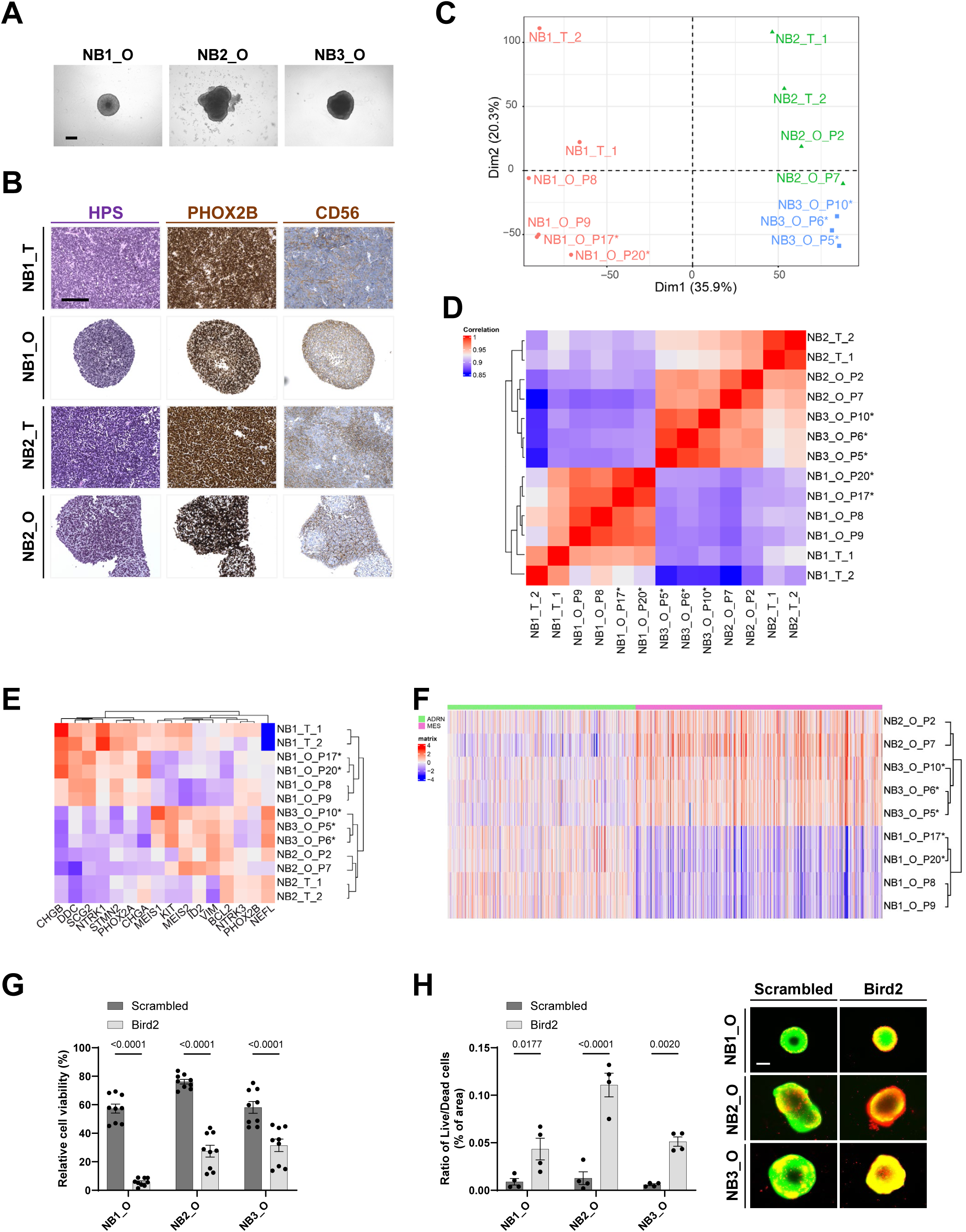
Targeting ITPR1-BCL2 interaction can kill *MYCN*-amplified neuroblastoma organoids. **A.** Representative bright-field images of neuroblastoma organoid lines 1, 2 and 3 after 10 days of culture. From day 0 to 10 post-seeding, neuroblastoma organoid lines expand to approximately 600 (NB1_O) and 800 μm (NB2_O and NB3_O) of diameter. Samples were obtained from patient-derived xenografts (NB1, NB2) and patient biopsy (NB3). Neuroblastoma organoid lines have been established and expanded using the protocol described in Methods. Scale bar: 300 μm. _O, organoid. **B**. Representative hematoxylin phloxine saffron (HPS) and immunohistochemistry characterization of neuroblastoma tumors and corresponding organoid lines using clinical markers routinely used for neuroblastoma diagnosis (PHOX2B, CD56). _T, tumor, _O, organoid. Scale bar: 150 μm. **C**. PCA of RNA-seq data from tumors (_T, 1 and 2 for replicates) and paired organoid lines (_O) plotted in 2D, using their projections onto the first two PCs (Dim1 and Dim2). Each data point represents one sample. _P, passage at time of collection; *, cryopreservation/revival. **D**. Heatmap of pairwise Pearson correlation coefficients based on global transcriptomic expression profile showing the clustering of neuroblastoma organoid lines (_O) with their paired tumor of origin (_T, 1 and 2 for replicates). **E**. Hierarchical clustering analysis based on the centered-normalized expression values of neuroblastoma tumors/paired neuroblastoma organoid lines (_O) and classical neuroblastoma differentiation markers (PMID: 18780787). _T, tumor; _P, passage at time of collection; *, cryopreservation/revival. **F**. Hierarchical clustering analysis based on the centered-normalized expression values of neuroblastoma organoid lines (_O) and mesenchymal (MES) and adrenergic (ADRN) signatures ^41^. _T, tumor; _P, passage at time of collection; *, cryopreservation/revival. **G**. Cell viability in neuroblastoma organoid (_O) lines 1, 2 and 3 monitored using CellTiter-Glo® assay at 72 h after treatment with Scrambled or Bird2 peptides. Scrambled peptide n=9, Bird2 peptide n=9. Mean +/- STDEV. U-test. P-values are indicated. **H**. Live/Dead assay in neuroblastoma organoid (_O) lines 1, 2 and 3. Quantification was performed by measuring the ratio of surface area of live to dead cells (left). Representative images of live (green)/dead (red) immunofluorescence stainings of neuroblastoma organoid treated for 72 h with Scrambled or Bird2 peptides. Scale bar: 200 μm. n = 4 spheres at least per condition. Representative results of two independent experiments. U-test. P-values are indicated.

Finally, the three *MYCN*-amplified NB_O lines were treated by the Bird2 or control peptide. We observed a strong decrease in cell viability associated with a rapid death of the tumoral cells in both neuroblastoma organoid models (Figure 8 G-H).

## DISCUSSION

MYC transcription factors are essential in promoting cell proliferation in different physiological and pathological contexts, including during cancer development. However, they also regulate cell death and cellular senescence, which counteract tumor formation, when highly activated in normal cells. In this study, we identified *ITPR1* as a new direct target gene of MYC and demonstrated its critical role in inducing ER-mitochondria Ca^2+^ transfer, cell death and cellular senescence. In line with these discoveries, *ITPR1* expression is low in many cancers and restoring MYC/ITPR1 signaling induces cancer cell death. Strikingly, we identified some cancer cells, with high expression of *MYC* or *MYCN* and of *BCL2*, displaying higher expression levels of *ITPR1*, such as *MYCN*-amplified neuroblastoma cells. In those aggressive tumors, NMYC controls *ITPR1* expression levels, high *ITPR1* expression correlated with a lower patient survival, and impeding the inhibition of ITPR1 activity by BCL2 using a blocking peptide killed those neuroblastoma cells and decreased neuroblastoma tumor volume *in vivo*.

MYC factors can directly control the expression of a wide range of genes involved in a broad spectrum of cellular effects. Among them, MYC controls the expression of genes promoting cell proliferation (e.g., cyclins, E2F) or cell cycle arrest (e.g., p15INK4B, p14ARF) as well as cell death (e.g., BIM, p14^ARF^). The expression of genes controlling the latter two cell fates is decreased during MYC-induced tumorigenesis ^1,4,7^. Other target genes, such as genes regulating cell metabolism (e.g., several glycolytic enzymes), are also critical regulators of MYC response, for instance to ensure energy production in line with the needs of dividing cells ^3,43^. Here, we identified *ITPR1* as a new direct target gene of MYC. ITPR1 is particularly interesting and quite unique in the target genes of MYC as it can participate in the different effects, eventually opposite, of MYC. Indeed, ITPR1, by controlling Ca^2+^ signaling, can impact various pathways and cell fates, including cell proliferation, cell death, cellular senescence and metabolism. For instance, Ca^2+^ is required for cell proliferation as it acts as a co-factor of many enzymes, including mitochondrial dehydrogenases whose activities are required for mitochondrial activity and ATP generation, whereas mitochondrial Ca^2+^ overload is sufficient to induce cell death or cellular senescence ^12–18,22,23^. Strongly supporting a key role of ITPR1 and mitochondrial Ca^2+^ accumulation in MYC-induced safeguard programs, MYC activation increases mitochondrial Ca^2+^ and reducing the level of ITPR1, involved in Ca^2+^ release from the ER, or VDAC3, involved in Ca^2+^ entry into the mitochondria, inhibits MYC-induced cell death and senescence.

As other tumor suppressors activated by MYC (p14^ARF^, p53, BIM…) ^27,44–46^, *ITPR1* was downregulated in most types of cancers, and increasing its activity promoted cancer cell death. Moreover, our data support a general decrease in ER-mitochondria Ca^2+^ transfer in cancer cells, to limit Ca^+^ accumulation and cell death, decreasing their sensitivity to MYC-induced safeguard mechanisms. They are also consistent with a more glycolytic phenotype of most cancer cells, given that Ca^2+^-dependent enzymes of the TCA (tricarboxylic acid) cycle would have a more limited access to Ca^2+ 16,47^.

Unexpectedly, we found molecular features of some cancer cells displaying high *ITPR1* expression levels, alongside high levels of expression of *MYCN* and/or *MYC*, and of *BCL2*, encoding an anti-apoptotic protein able to inhibit ITPR1 Ca^2+^ channel activity ^30^. This is particularly the case in *MYCN*-amplified neuroblastoma. Neuroblastomas are childhood tumors accounting for 5-10% of all pediatric tumors, causing around 15% of cancer-associated childhood deaths. High-risk neuroblastomas have a poor outcome with a long-term survival around 50%. Among these, 20% show amplification of the *MYCN* gene and another 10% display a high *MYC* expression. These two categories are part of the most aggressive neuroblastomas ^10,48^. We showed that higher expression levels of *ITPR1* in *MYCN*-amplified neuroblastomas were correlated with an increased risk of death. Mechanistically, we speculate that patients with *MYCN* amplification and high *ITPR1* expression levels have a very poor prognosis, because of their highest levels of NMYC and/or BCL2, well-known factors promoting tumor resistance and/or progression. These patients with the worse prognosis may benefit from a new therapeutic strategy impeding the inhibition of ITPR1 activity by BCL2. Indeed, the Bird2 peptide, known to block the ITPR1/BCL2 interaction and to increase mitochondrial Ca^2+ 31^, promoted the death of all *MYCN*-amplified neuroblastoma cell lines and patient-derived organoids tested, and decreased the size of neuroblastoma tumors in chick embryos. Neuroblastomas arise in the course of the sympathoadrenal lineage differentiation and is associated with a high degree of tumor cell plasticity, intratumor and interpatient heterogeneities ^49^. Impeding the inhibition of ITPR1 activity by BCL2 could help designing more effective treatment regimens for high-risk neuroblastoma.

Overall, our study demonstrates (i) that TFs of the MYC family orchestrate ER-mitochondria Ca^2+^ transfer by activating the transcription of the *ITPR1* gene to induce cell death and senescence, (ii) that this pathway is generally inhibited in cancers, and (iii) that when it can be reactivated, such as in *MYCN*-amplified neuroblastomas, it constitutes a novel vulnerability for aggressive tumors. Our study thus unveils an exciting new field of research as it paves the way for future investigations on the role of MYC-induced Ca^2+^ signaling in controlling other MYC-dependent molecular, cellular and organismal phenotypes, in particular in the context of aging and cancer, and for the potential development of novel therapeutic approaches.

## MATERIALS AND METHODS

### Cell culture and reagents

MRC5 normal human embryonic lung fibroblasts, U2OS cancer cells (ATCC, Manassas, VA, USA) and virus-producing cells 293-GP and 293-T (Clontech) were cultured in Dulbecco′s modified Eagle′s medium (DMEM, Life Technologies) with GlutaMax supplemented with 10% fetal bovine serum (FBS) (Life Technologies) and 1% penicillin/streptomycin (Life Technologies). Kelly (Sigma-Aldrich), KPNYN (NIBIOHN JCRB), (CLB BERLud 1, CLB BAR Rec, CLB-Ga (Centre Léon Bérard) neuroblastoma cells were cultured in RPMI 1640 medium (Life Technologies) with 10% FBS (Sigma) except for Kelly which were cultured at 20% FBS (Life Technologies), and 1% penicillin/streptomycin (Life Technologies). SKNAS, SKNSH, SHSY5Y neuroblastoma cells (Sigma-Aldrich) were cultured in DMEM with 10% FBS and 1% penicillin/streptomycin. All cells were grown in standard conditions (37°C, 5% CO_2_). Experiments were performed on Mycoplasma-negative cells.

MRC5-MYC:ER cells were treated with 100 nM (Z)-4-hydroxytamoxifen (4OHT, H7904, Sigma-Aldrich) to activate MYC. Kelly cells were treated with 5 µM VPC-70619 (HY-144878, MedChem Express) to inhibit NMYC. In 2D experiments, neuroblastoma derived cells were treated with 20 µM of Scrambled (sequence: RKKRRQRRRGGDLNEVTCSLIVDRINPVKLY) or Bird2 (sequence: RKKRRQRRRGGNVYTEIKCNSLLPLAAIVRV) peptides (Smart Bioscience).

### Plasmids, plasmid transfection and infection

Retroviral vectors, pBabe-puro-MYC:ER ^50^, encoding MYC oncogene fused to the ligand binding domain of the estrogen receptor (ER) (Addgene plasmid #19128), pBabe-puro (Addgene plasmid #1764) ^51^, pLNCX2-mito-GEM-GECO1, to measure mitochondrial Ca^2+ 21^, were used in MRC5 cells. Lentiviral vectors pLKO.1 encoding shRNAs targeting MYCN (TRCN0000020695 for shMYCN.1 and TRCN0000020697 for shMYCN.2, Sigma-Aldrich) were used to knock down MYCN and pMOS028:Mitycam lentiviral vector (Addgene plasmid #163046) ^52^ was used to measure mitochondrial Ca^2+^ in Kelly cells. 293-GP (for retrovirus production) or 293-T (for lentivirus production) were transfected with these plasmids in OptiMEM medium (Gibco) using PEIpro transfection reagent (Polyplus) according to the manufacturer’s recommendations. 48 h after transfection, viral supernatants were collected, diluted in DMEM (for MRC5 or U2OS cells) or RPMI 1640 (for Kelly cells) and hexadimethrine bromide (8 μg/mL, Sigma-Aldrich) was added. Viral supernatant was added to MRC5, U2OS or Kelly cells, centrifuged at 2,000 RPM for 30 min and then incubated for 8 h. 24 h after infection, selection was started using puromycin (InvivoGen) at 500 ng/mL or geneticin (Life Technologies) at 100 μg/mL.

### siRNA transfection

ON-TARGETplus siRNA SMARTpools of 4 siRNAs (Horizon Discovery) targeting human ITPR1, ITPR2, ITPR3, MYC, MAX, SUPT5H, TRRAP, KAT2A (GCN5), KAT5 (TIP60), RUVBL1 (TIP49) or RUVBL2 (TIP48) or VDAC3 were used, or ON-TARGETplus non-targeting pool as negative control (siControl) (Horizon Discovery). siRNAs were incubated for 20 min with DharmaFECT 1 transfection reagent (Horizon Discovery) in antibiotics and serum-free DMEM. MRC5, MRC5-MYC:ER or U2OS-MYC:ER cells were then reverse transfected with this mix in antibiotic-free DMEM containing 10% FBS. Final concentration of siRNAs was 15 nM. The next day, medium was changed with DMEM containing 1% antibiotics and 10% FBS.

### RNA extraction, reverse transcription and real-time quantitative PCR

RNAs were isolated using NucleoZol (Macherey-Nagel) following the manufacturer’s recommendations, quantified by NanoDrop One C (Thermo Fisher Scientific) and reverse-transcribed to cDNA with Maxima First Strand cDNA Synthesis Kit (Thermo Fisher Scientific) according to the manufacturer’s instructions. Real-time quantitative PCR were performed with ONEGreen FAST qPCR Premix (Ozyme) following the manufacturer’s recommendations, on a Bio-Rad CFX96 system. Primer sequences are listed in Table S3. mRNA levels of housekeeping genes GAPDH and PGK1 were used for normalization in MRC5 cells, GAPDH in U2OS cells and TBP in Kelly neuroblastoma cells. Relative mRNA levels were calculated using the comparative Ct (ΔΔCT) method.

### Western blot

After washing with 1X PBS, cells were scraped in Laemmli buffer containing 10% SDS, 10% glycerol and 1 M TrisHCl pH 6.8 and proteins in cell lysates were quantified using NanoDrop One C (Thermo Fisher Scientific). Bromophenol Blue and β-mercaptoethanol were added and samples were boiled at 95℃ for 10 min. Proteins were separated on SDS-PAGE in TG-SDS migration buffer (Euromedex) and transferred to nitrocellulose membranes (Bio-Rad). Membranes were then blocked with 5% milk in TBS with 0.01% Tween-20 (Sigma-Aldrich) TBST for 1 h at room temperature and incubated with mouse primary antibodies against ITPR1 (sc-271197, Santa Cruz Biotechnology) or α-Tubulin (T6199, Sigma-Aldrich) at 4℃ overnight with gentle shaking. After removing primary antibodies, membranes were washed three times for 10 min with TBST and incubated with anti-mouse secondary antibody conjugated to horseradish peroxidase (715-035-150, Jackson ImmunoResearch) for 1 h at room temperature, and again washed with TBST. Chemiluminescence was revealed with Clarity MAX Western ECL Substrate (Bio-Rad) for ITPR1 and Pierce ECL Western Blotting Substrate (Thermo Fisher Scientific) for α-Tubulin, using ChemiDoc XRS (Bio-Rad) and Image Lab software.

### Crystal violet staining, cell death analysis and senescence-associated-β-galactosidase assay

For crystal violet staining, cells were washed once with 1X PBS, fixed for 10 min with 3.7% formaldehyde (Sigma-Aldrich) and then stained with crystal violet solution (Sigma-Aldrich). For cell death analysis, the relative number of dead cells was either counted manually following cell incubation with trypan blue (Life Technologies) or monitored by Incucyte live-cell imaging system (Sartorius) following cell incubation with SYTOX Green (Thermo Fisher Scientific). Senescence-associated-β-galactosidase (SA-β-gal) assay was performed as previously described in ^53^. The percentage of SA-β-gal-positive cells was calculated after counting at least 100 cells per condition.

### Mitochondrial calcium measurement

MRC5-MYC:ER cells expressing mito-GEM-GECO1, a mitochondrial calcium ratiometric gene reporter, were transfected with siRNAs and treated with 4OHT in 96-well plates. Two days after 4OHT treatment, cells were washed twice with 1X PBS and then incubated 15 min in HBSS with Ca^2+^ and Mg^2+^ (Life Technologies) before image acquisition. Images were acquired on a Zeiss LSM 980 confocal microscope, with one excitation performed at 405 nm and two emissions recorded at 437-499 nm and 520-755 nm. Pictures were analyzed with ImageJ-Fiji software. In each cell, fluorescence intensity was measured in three different areas. The ratio F(437-499)/F(520-755) was calculated for each area and the mean of the ratios of the three areas was calculated for each cell.

Kelly cells expressing Mitycam mitochondrial calcium genetic sensor were seeded in 96-well plates in RPMI 1640 medium without red phenol. Scrambled or Bird2 peptides diluted in HBSS with Ca^2+^ and Mg^2+^ were added to the wells the following day. Fluorescence was recorded 50 s before peptide injection and during 5 min in total with a picture taken every 3 s. Images were acquired using Opera Phenix HCS (Perkin Elmer), with excitation at 513nm and emission recorded at 530nm and the Harmony software was used to analyze them. Fluorescence intensity was measured in three different areas per cell and the ratio (F1-F0)/F0 (F1 being the maximum fluorescence intensity after adding peptides and F0 the fluorescence intensity before adding peptides) was calculated for each area and the mean ratio of the three areas was calculated for each cell.

### Chick embryo model of neuroblastoma

Oncofactory SAS (an ERBC company) performed the experiments following the AVI-cellDXTM procedure described in ^36,37,54^. Shortly 2500 Kelly-GFP cells were injected into the dorsal roof of the neural tube, in HH14 stage (E2) embryos, at the level of somites 18 to 24. The day after (E3), 10 mg/kg of control Scrambled peptide or of Bird2 peptide were intravenously injected. At HH25 (E4), embryos were harvested, weighted and measured, and a quantitative analysis of the tumor volume in each embryo was performed using 3D light sheet microscopy (Miltenyi Biotec) and Imaris software. The results are presented as a normalized tumor volume to the body surface area (BSA, calculated with the Dubois&Dubois formula).

### Neuroblastoma fresh tissue collection

Patient-derived xenograft models were provided by the St. Jude Children’s Research Hospital. Implantation was done according to their guidelines ^55^. NSG-NOD SCID mice were obtained from Charles River animal facility. The mice were housed in sterilized filter-topped cages and maintained in the P-PAC pathogen-free animal facility (D 69 388 0202). All animal studies were performed in strict compliance with relevant guidelines validated by the local Animal Ethic Evaluation Committee (C2EA-15) and authorized by the French Ministry of Education and Research (Authorization APAFIS#28836). Human tissue sample was obtained through a biopsy performed at Centre Léon Bérard. This sample was collected in the context of patient diagnosis. The Biological Resource Centre (BRC) of the Centre Léon Bérard (n°BB-0033-00050) and the biological material collection and retention activity are declared to the Ministry of Research (DC-2008-99 and AC-2019-3426). The study had all necessary regulatory approvals and informed consents are available for all patients.

### Derivation and culture of neuroblastoma organoids

Fresh tissues were minced into small pieces and digested with collagenase D (0.125 mg/mL Roche) and 1 µg/mL DNase I (Sigma) diluted in HBSS (Gibco). After 90 min incubation at 37 °C cells were washed using advanced DMEM/F-12 medium (Gibco). Then, cultures were established in 96-well ULA plates (Corning, cat. no. 7007) in a culture medium, which consists of advanced DMEM/F-12 medium (Gibco), 1X B-27 supplement without vitamin A (Gibco), 1X N2 supplement (Life), 40 ng/mL hFGF-b (Peprotech), 20 ng/mL hEGF (Peprotech), 10 ng/mL hPDGF-AA (Peprotech), 10 ng/mL hPDGF-BB (Peprotech) and 6000 U/mL heparin (Sigma). Medium was changed twice a week and neuroblastoma organoids were split every 10 days when reaching a diameter of 600-800 µm using TrypLE Express Enzyme (ThermoFisher Scientific). All cultures were tested monthly for mycoplasma using the MycoAlert® Mycoplasma Detection Kit (Lonza), in accordance with the manufacturer’s instructions.

### Histological analyses

Tissues and neuroblastoma organoids were fixed and processed as described before ^56^ in close collaboration with the Research pathology platform East (Anapath Recherche, Cancer Research Center of Lyon (CRCL), Lyon, France. In brief, 4 µm slides were stained with HPS (Hematoxylin Phloxine Saffron) or the following antibodies: PHOX2B (1/200, FNab06409, FineTest), CD56 (1/100, NCL-L-CD56-504, Leica biosystems) and then incubated with relevant antibody-HRP conjugates for 1 h at room temperature (RT), revealed with 3,3′-diaminobenzidine (DAB) for 5 min, and counterstained with Gill’s-hematoxylin. Slides were mounted using Pertex (Histolab) and observed with EVOS™ M7000 Imaging System.

### CellTiter-Glo® cell viability and LIVE/DEAD™ assays

Neuroblastoma organoids were dissociated with TrypLE Express (Thermo Fisher Scientific, cat. no. 12605036) and seeded at 10.000 cells/well in 96-well plates (Corning, cat. no. 4515). They were allowed to form for 4 days and were then treated with vehicle (DMSO), Scrambled or Bird2 peptides at 10 µM. CellTiter-Glo® cell viability (Promega, cat. no. G9683) and LIVE/DEAD™ (Thermo Fisher Scientific, cat. no. 10237012) assays were performed after 72 h of treatment in accordance with the manufacturer’s instructions. For cell viability assay data variation was expressed as mean ± SD. For LIVE/DEAD assay neuroblastoma organoids were imaged using EVOS™ M7000 Imaging System and LIVE/DEAD stainings were quantified by measuring the surface area of live and dead cells on at least four neuroblastoma organoids on 2 independent experiments using ImageJ software.

### RNA sequencing

1000 ng of total RNAs from tissues and neuroblastoma organoids were isolated using AllPrep kit (Qiagen, Cat. No. 80204) and following the manufacturer’s instructions. Libraries were prepared with Illumina Stranded mRNA Prep following recommendations and sequenced using a NovaSeq 6000 according to the standard Illumina protocol by the Cancer Genomics Platform from the CRCL. Quality control of reads was performed using FastQC. Reads were mapped to the GRCh38 human genome using STAR (v 2.7.10a) and Ensembl annotations (v 108). Gene counts were then computed using HTseq-count (v 2.0.2) and loaded in R (v 4.3.3) and converted into a DESeq2 (v 1.42.0) object. Expression was normalized with vst. Heatmaps were made using the package ComplexHeatmap (v 2.18.0). FactoMineR (v 2.10) and factoextra (v 1.0.7) were used for the Principal Component Analysis (PCA) and graphical representation, respectively.

### Bioinformatics analysis

All genomic data were analyzed with R/Bioconductor packages, R version 4.2.2 (2022-11-10) [https://cran.r-project.org/; http://www.bioconductor.org/] in a linux environment (x86_64-pc-linux-gnu [64-bit]). ***ChIP seq.*** MYC binding on the *ITPR1* promoter was analyzed using publicly available ChIPseq data. The following datasets were analyzed: GSE44672, GSE86412, GSE80154, GSE80151. ChiPseq peak chromosomal locations were downloaded into R and plotted using karyoploteR package. When available ChiPseq occupancy data (i.e., wiggle and bigwig) were also downloaded and plotted. Results were presented using coverage plots, described briefly. Coverage plots: Bigwig files for each sample were used to extract the ChIPseq signal corresponding to each of the selected regions. The context given by the bottom tracks includes the chromosomal location, the annotated genes, and the “cCRE” ENCODE track. This track summarizes the ENCODE Candidate Cis-Regulatory Elements (cCREs) combined from all cell types, as described here: http://genome-euro.ucsc.edu/cgi-bin/hgTrackUi?hgsid=290804634_bByAayidA2MzaofYyPM9j7hRtFmU&db=mm10&c=chr9&g=encodeCcreCombined.

### RNAseq

TCGA genomic data were analyzed with R/Bioconductor packages and associated packages (TCGAbiolinks, edgeR, ggplot2, singscore, msigdbr, and clusterProfiler). RNAseq data were retrieved from TCGA PanCancer and GSE80154 dataset (only focused on neuroblastoma).

### Survival analysis

To perform survival analysis, the dataset GSE49710 was used. It included survival information for 498 neuroblastoma patients. Kaplan-Meier survival curves were generated using survival analysis packages (survival and survminer). Log-rank tests were applied to assess differences in survival between groups. Cox proportional hazards regression models were employed to evaluate the impact of variables on survival. Stratification was performed based on MYCN amplification. Significance was set at p < 0.05.

### Statistical analysis

GraphPad Prism 9 was used to perform statistical analysis and create graphs, which are presented as mean of three or more independent experiments with SEM, except stated otherwise in figure legend. Statistical tests used are indicated in figure legends and p-values are indicated in each graph.

## DATA AVAILABILITY

The source data are available upon request to the corresponding authors.

## AUTHOR CONTRIBUTIONS

XM, AT, MV, KZ, JEH, DZ, AH, GR, CM, CC, FB, VA, MC, OG, CK, IA, JCH, FJ, HHV, LB and NM performed experiments, provided key materials and/or drew the figures. XM, AT, MV, KZ, BM, CV, BD, CDB, LB, NM and DB designed the experiments and the results were analyzed by all the co-authors. LB designed, supervised and got funding for the organoids work. NM and DB designed the overall study, co-supervised the work and wrote the manuscript with input from all authors.

## ACKNOWLEDGEMENTS

We thank Brigitte Manship for critical reading of the manuscript. We thank Marie Castets, Geert Bultynck, Gabriel Ichim and Benjamin Gibert for sharing tools, expertise and/or advices. We thank the patients and their families who consented to participate in this study and the BRC of CLB and HCL (Tissu-Tumorothèque Est). We thank the “Mia Neri foundation” that support this work. We thank the St. Jude Children’s Research Hospital as the provider of the neuroblastoma patient-derived xenografts. We thank the platforms participating to this work: Cell Imaging Platform at the CRCL, the Centre d’Imagerie Quantitative Lyon-Est (SFR Centre d’Imagerie Quantitative Lyon-Est), the Research pathology platform East at CRCL and the Cancer Genomics Platform from the CRCL. This study was funded by grants from Fondation ARC to NM; la Ligue Nationale contre le Cancer, Association Hubert-Gouin – Enfance & Cancer, Association Enfants Cancers Santé and la Société Française de lutte contre les Cancers et les leucémies de l’Enfant et de l’adolescent, INSERM Transfert to DB. KZ was supported by China Scholarship Council (202008340070). MV was supported by the Fondation de France and CM was supported by la Ligue Nationale contre le Cancer. This research was supported by the Foundation ARCECI innovation award (USA) and the ANR JCJC program (CHILD-SARC; ANR-19-CE14-0001-01) awarded to LB. FB received financial support from the “Mia Neri foundation”. CC received financial support from Bristol Myers Squibb (BMS). CK and OG were supported by a grant from the Comités de l’Ardèche et du Rhône de la Ligue contre le cancer. CK and OG wish to thank Sandrine Gonin-Giraud for sharing expertise and advices.

