## Supplemental figures and tables for "MYC shapes ER-mitochondria calcium transfer by directly targeting *ITPR1*: implications for MYC-induced safeguard mechanisms and cancer"

**SUPPLEMENTARY FIGURE LEGENDS**

**Supplementary Figure 1. A.** RT-qPCR of *MYC*, *ITPR1*, *ITPR2* and *ITPR3* genes at day 3 after transfection of MRC5 cells with a control siRNA pool (siControl) or with a siRNA pool targeting MYC (siMYC). Mean +/- SEM of n = 4 independent experiments. Two-Way ANOVA. P-values are indicated. **B.** Representation of MYC occupancy at the *Itpr1* promoter in mouse cells: mouse embryonic fibroblasts (MEFs) and T cells (GSE44672). **C-D**. RT-qPCR of *ITPR1* and *MAX* genes in MRC5-MYC:ER cells transfected with siControl or with a siRNA pool targeting MAX (siMAX) (**C**) or RT-qPCR of *ITPR1* and *TRRAP* genes in MRC5-MYC:ER cells transfected with siControl or with a siRNA pool targeting TRRAP (siTRRAP) (**D**), and then treated or not with 4OHT for 3 days. Mean +/- SEM of n = 4 independent experiments. Two-Way ANOVA. P-values are indicated.

**Supplementary Figure 2.** MRC5-MYC:ER cells were transfected with a control siRNA pool (siControl) or with the indicated siRNA pool and then treated or not with 4OHT. **A.** Crystal violet staining at day 10 after 4OHT treatment. Representative image of n = 3 independent experiments. **B**. Representative images (upper panel) and quantification (lower panel) of SA-β-galactosidase-positive cells at day 3 after 4OHT treatment. Mean +/- SEM of n=3 independent experiments. Two-Way ANOVA. P-values are indicated.

**Supplementary Figure 3.** MRC5-MYC:ER cells were transfected with a control siRNA pool (siControl) or with a siRNA pool targeting ITPR1 (siITPR1) or VDAC3 (siVDAC3) and then treated or not with 4OHT. Representative images (left) and quantification (right) of SA-β-galactosidase-positive cells at day 3 after 4OHT treatment. Mean +/- SEM of n = 3 independent experiments. Two-Way ANOVA. P-values are indicated.

**Supplementary Figure 4**. Expression data of *BCL2* and *ITPR1* extracted from a human neuroblastoma dataset (Tumor Neuroblastoma – SEQC – 498 – custom – ad44kcwolf) using the R2 Genomics Analysis and Visualization Platform. Correlation of expression between *BCL2* and *ITPR1* mRNA levels is shown in 401 *MYCN*-non-amplified neuroblastomas.

**Supplementary Figure 5. A**. RT-qPCR of *MYCN* and *ITPR1* genes in Kelly cells expressing a control non-targeting shRNA (shNEG) or shRNAs targeting MYCN (shMYCN). Mean +/- SEM of n = 3 independent experiments. Two-Way ANOVA. P-values are indicated. **B.** RNA-seq validation of *MYCN* expression level upon its depletion in SHEP21 using a TET-OFF system (GSE80154). Wilcoxon test. P-values are shown.

**Supplementary Figure 6. A.** Kaplan-Meier survival curves drawn from GSE49710 dataset from patients with *MYCN*-non-amplified neuroblastoma. The 2 groups were formed according to the lowest quartile of *ITPR1* expression. **B**. Representative images of Kelly and SKNAS neuroblastoma cells after 24 h treatment with Scrambled or Bird2 peptides. Cell death was monitored using SYTOX Green.

### Supplementary Figure 1

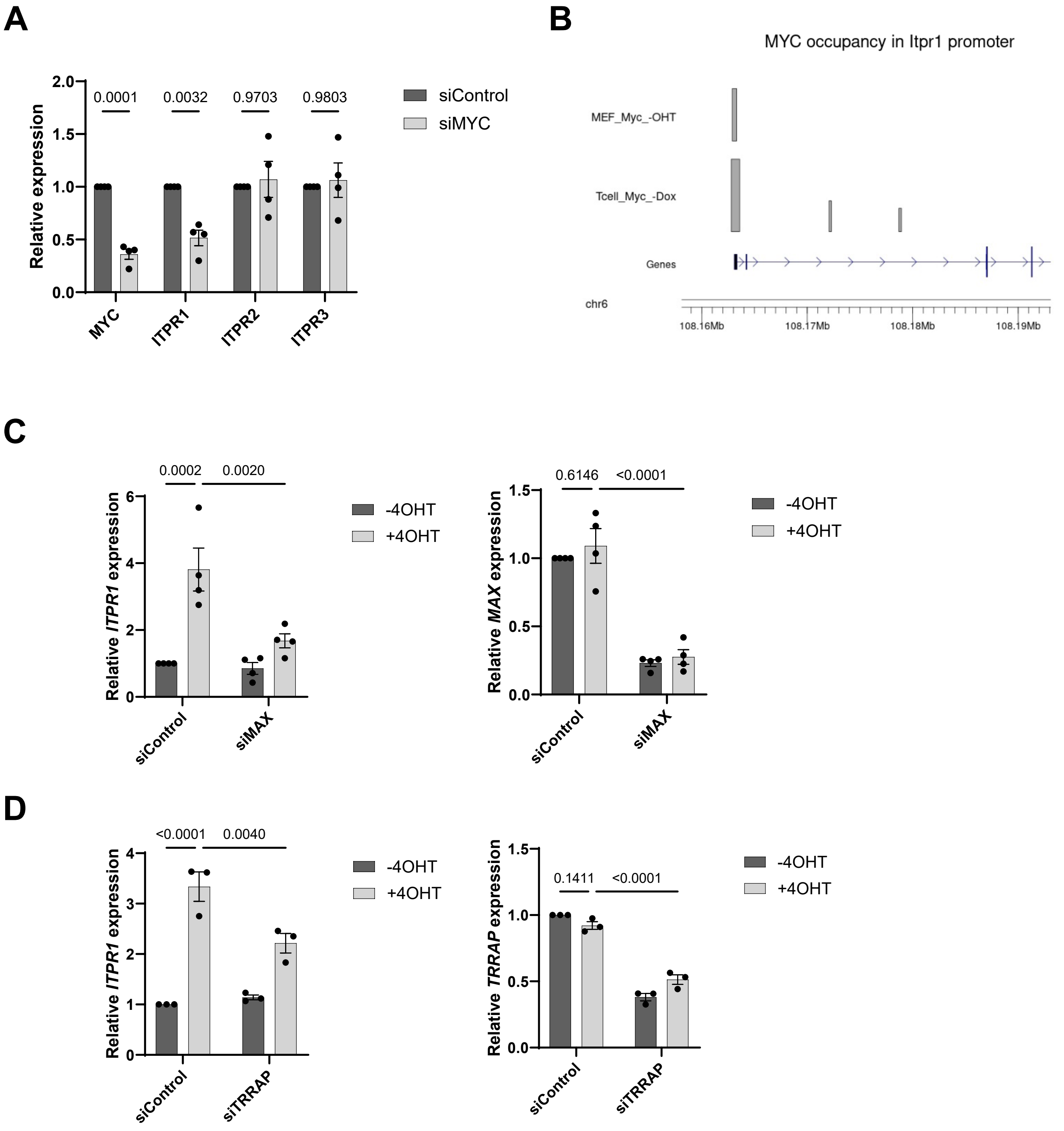

#### Supplementary Figure 2

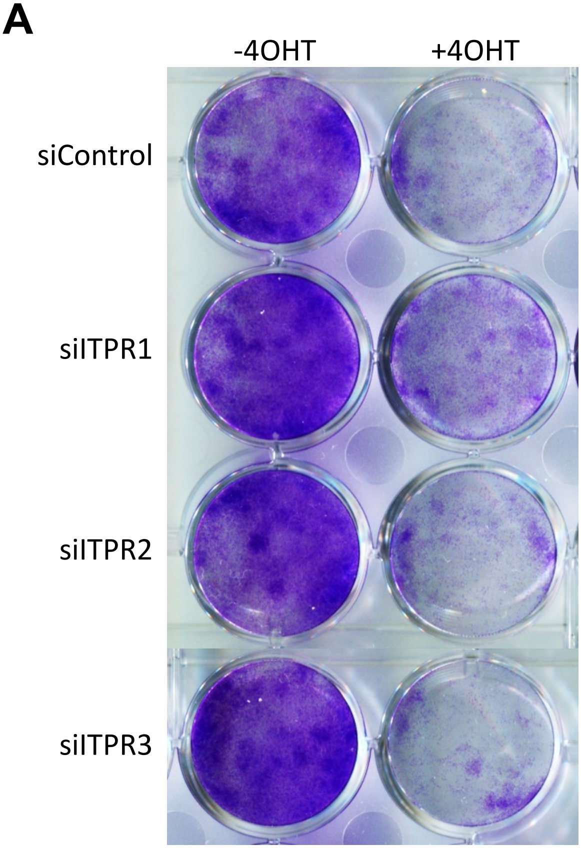


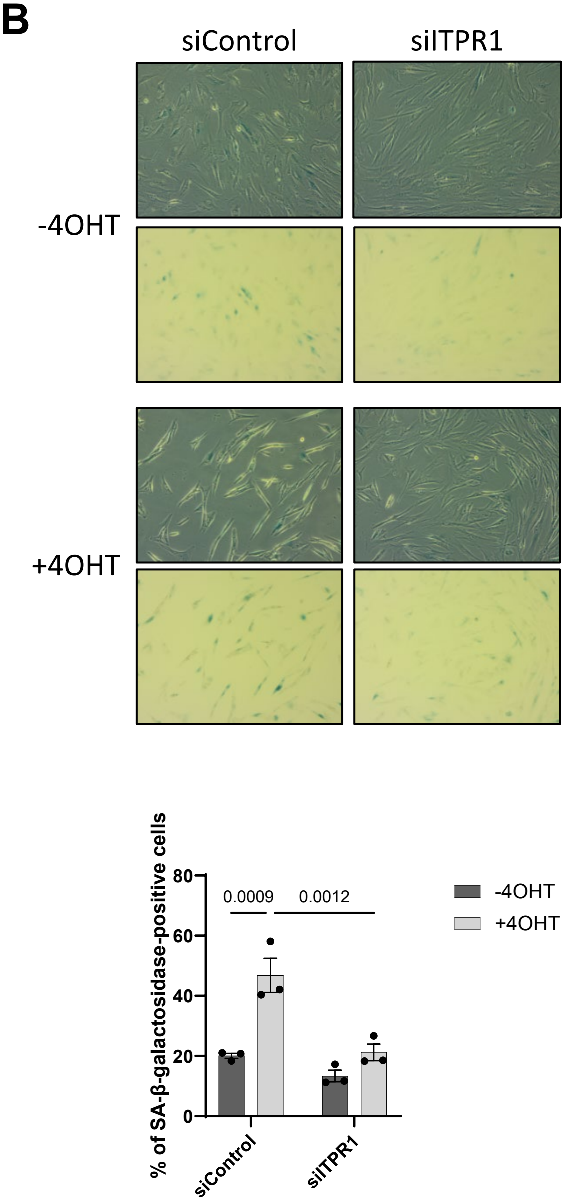

### Supplementary Figure 3

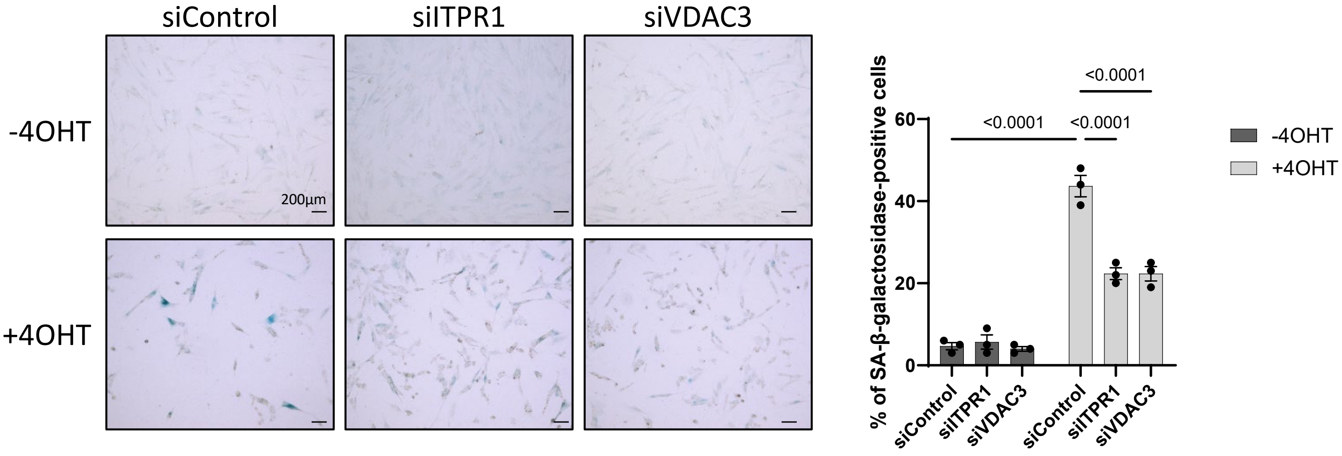

### Supplementary Figure 4

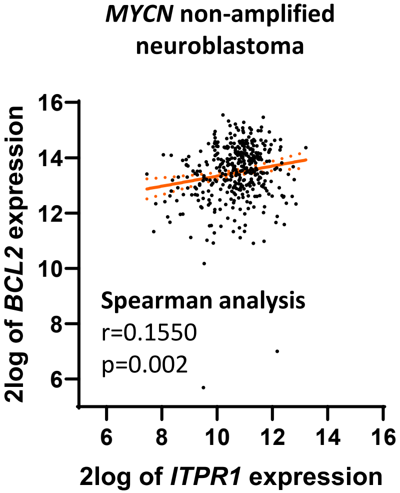

### Supplementary Figure 5

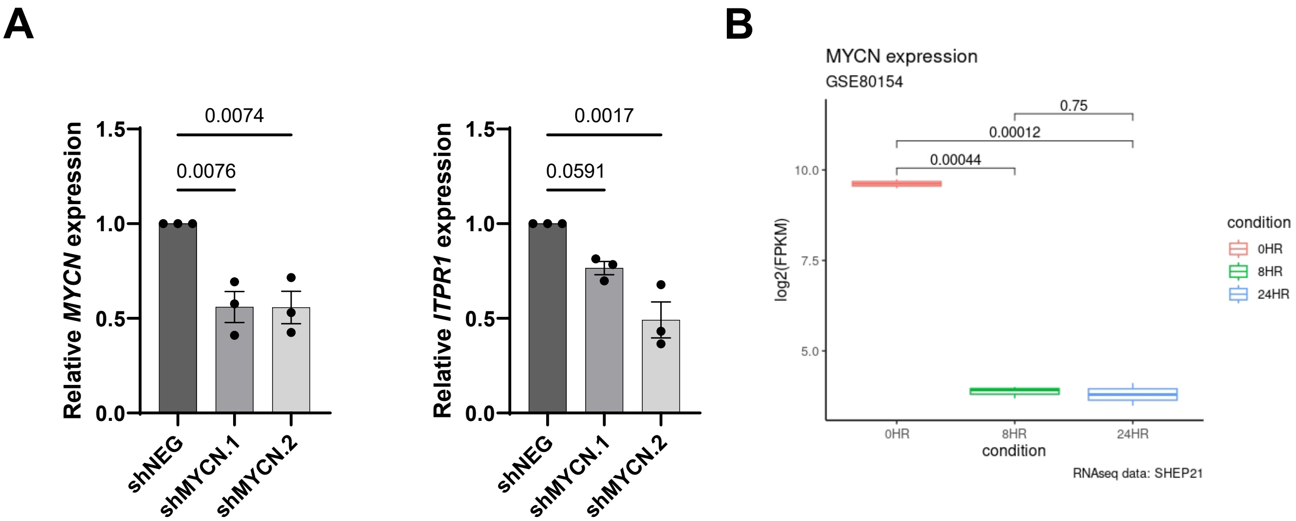

### Supplementary Figure 6

A

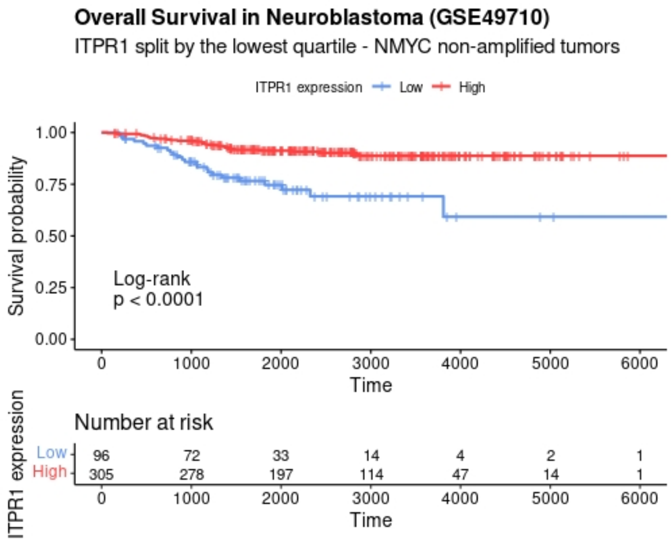


B

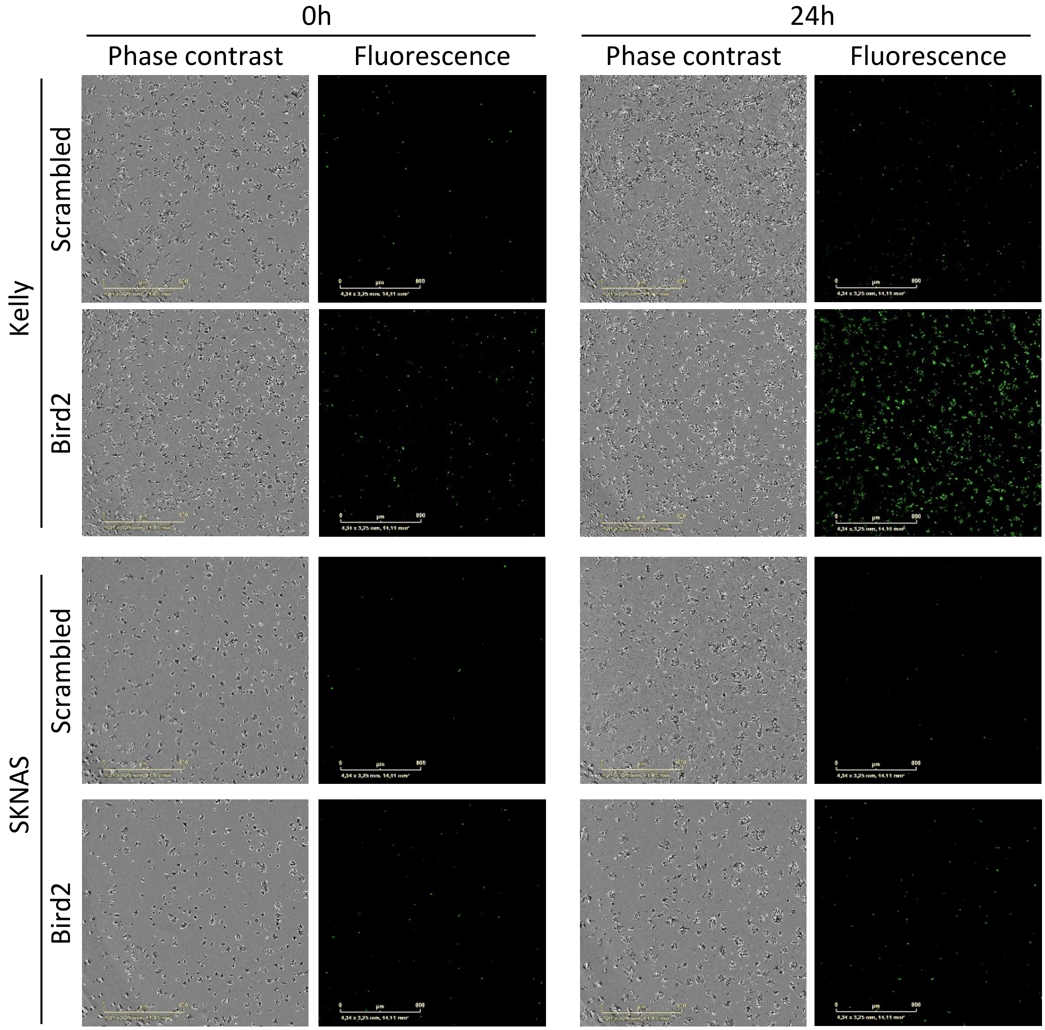

#### SUPPLEMENTARY TABLES

**Supplementary Table 1.** Contribution of different MYC partners and co-factors in the induction of *ITPR1* expression by MYC. MRC5-MYC:ER cells were transfected with siControl or with siRNA pools targeting MYC partners and co-factors listed in the table and were then treated or not with 4OHT for 3 days. RT-qPCR for checking *ITPR1* gene expression and knockdown efficiency were performed. Mean +/- SEM of n=3 independent experiments were calculated and Two-Way ANOVA test was performed. P-values for the reversal of *ITPR1* induction by MYC are indicated.

| **Gene targeted by siRNA** | **P-value for the reversal of *ITPR1* induction by MYC** |
|---|---|
| KAT2A | 0.330 |
| KAT5 | 0.414 |
| MAX | 0.002 |
| RUVBL1 | 0.139 |
| RUVBL2 | 0.559 |
| SUPT5H | 0.516 |
| TRRAP | 0.004 |

**Supplementary Table 2.** List of the 54 cancer cell lines with high expression levels of *ITPR1* and *BCL2*. Name of cell lines, tissue of origin and type of cancer are indicated.

| ***ITPR1* and *BCL2* high cancer cell lines** | | |
|---|---|---|
| **Cell lines** | **Tissue of origin** | **Type of cancer** |
| KPNYN | Autonomic ganglia | Neuroblastoma |
| KELLY | Autonomic ganglia | Neuroblastoma |
| CHP126 | Autonomic ganglia | Neuroblastoma |
| NH6 | Autonomic ganglia | Neuroblastoma |
| KPNRTBM1 | Autonomic ganglia | Neuroblastoma |
| L540 | Haematopoietic and lympoid tissue | Hodgkin Lymphoma |
| HDLM2 | Haematopoietic and lympoid tissue | Hodgkin Lymphoma |
| JVM2 | Haematopoietic and lympoid tissue | Mantle Cell Lymphoma |
| JVM3 | Haematopoietic and lympoid tissue | B-Cell Prolymphocytic Leukemia |
| EB1 | Haematopoietic and lympoid tissue | Burkitt Lymphoma |
| OCIAML2 | Haematopoietic and lympoid tissue | Acute Myeloid Leukemia |
| THP1 | Haematopoietic and lympoid tissue | Acute Myeloid Leukemia |
| KCL22 | Haematopoietic and lympoid tissue | Chronic Myeloid Leukemia, BCR-ABL1+ |
| MUTZ5 | Haematopoietic and lympoid tissue | B-Lymphoblastic Leukemia/Lymphoma |
| GDM1 | Haematopoietic and lympoid tissue | Acute Myeloid Leukemia |
| MUTZ3 | Haematopoietic and lympoid tissue | Acute Myeloid Leukemia |
| KG1 | Haematopoietic and lympoid tissue | Acute Myeloid Leukemia |
| L1236 | Haematopoietic and lympoid tissue | Hodgkin Lymphoma |
| WSUDLCL2 | Haematopoietic and lympoid tissue | Diffuse Large B-Cell Lymphoma, NOS |
| HUT102 | Haematopoietic and lympoid tissue | Mycosis Fungoides |
| MOLT16 | Haematopoietic and lympoid tissue | T-Lymphoblastic Leukemia/Lymphoma |
| NUDHL1 | Haematopoietic and lympoid tissue | Diffuse Large B-Cell Lymphoma, NOS |
| KE97 | Haematopoietic and lympoid tissue | Plasma Cell Myeloma |
| PL21 | Haematopoietic and lympoid tissue | Acute Myeloid Leukemia |
| P31FUJ | Haematopoietic and lympoid tissue | Acute Myeloid Leukemia |
| ME1 | Haematopoietic and lympoid tissue | Acute Myeloid Leukemia |
| F36P | Haematopoietic and lympoid tissue | Myelodysplastic Syndromes |
| KMS27 | Haematopoietic and lympoid tissue | Plasma Cell Myeloma |
| MONOMAC6 | Haematopoietic and lympoid tissue | Acute Monoblastic/Monocytic Leukemia |
| OCIAML5 | Haematopoietic and lympoid tissue | Acute Myeloid Leukemia |
| HS611T | Haematopoietic and lympoid tissue | Hodgkin Lymphoma |
| EHEB | Haematopoietic and lympoid tissue | B-Lymphoblastic Leukemia/Lymphoma |
| LOUCY | Haematopoietic and lympoid tissue | Adult T-Cell Leukemia/Lymphoma |
| EOL1 | Haematopoietic and lympoid tissue | Chronic Eosinophilic Leukemia, NOS |
| HNT34 | Haematopoietic and lympoid tissue | Acute Myeloid Leukemia |
| SIGM5 | Haematopoietic and lympoid tissue | Acute Monoblastic/Monocytic Leukemia |
| NCO2 | Haematopoietic and lympoid tissue | Chronic Myeloid Leukemia, BCR-ABL1+ |
| MOLM6 | Haematopoietic and lympoid tissue | Chronic Myeloid Leukemia, BCR-ABL1+ |
| MONOMAC1 | Haematopoietic and lympoid tissue | Acute Monoblastic/Monocytic Leukemia |
| HUNS1 | Haematopoietic and lympoid tissue | Plasma Cell Myeloma |
| SKMM2 | Haematopoietic and lympoid tissue | Plasma Cell Myeloma |
| MEC1 | Haematopoietic and lympoid tissue | Chronic Lymphocytic Leukemia/Small Lymphocytic Lymphoma |
| SUPB15 | Haematopoietic and lympoid tissue | B-Lymphoblastic Leukemia/Lymphoma |
| OCIMY5 | Haematopoietic and lympoid tissue | Plasma Cell Myeloma |
| MOLM13 | Haematopoietic and lympoid tissue | Acute Myeloid Leukemia |
| KASUMI1 | Haematopoietic and lympoid tissue | Acute Myeloid Leukemia |
| AMO1 | Haematopoietic and lympoid tissue | Plasma Cell Myeloma |
| SKM1 | Haematopoietic and lympoid tissue | Acute Myeloid Leukemia |
| GRANTA519 | Haematopoietic and lympoid tissue | Mantle Cell Lymphoma |

| | | |
|---|---|---|
| KO52 | Haematopoietic and lympoid tissue | Acute Myeloid Leukemia |
| PLB985 | Haematopoietic and lympoid tissue | Acute Myeloid Leukemia |
| KASUMI6 | Haematopoietic and lympoid tissue | Acute Myeloid Leukemia |
| TUHR14TKB | Kindney | Renal Cell Carcinoma |
| NCIH2172 | Lung | Non-Small Cell Lung Cancer |

**Supplementary Table 3.** Sequences of primers used for RT-qPCR.

| Gene | Primer 1 | Primer 2 |
|---|---|---|
| ITPR1 | TACCCAGCGGCTGCTAAC | TGCAAATCCTGCTCCTCTGT |
| ITPR2 | AAAGCCTCAGTGGAATCCTGT | ATGGCAATTCCACGATTTTT |
| ITPR3 | CTGCTGTAGCCAGTGCAGAC | GGAGCAAGATCGTCCATCA |
| MYC | GCTGCTTAGACGCTGGATTT | TAACGTTGAGGGGCATCG |
| MYCN | CCACAAGGCCCTCAGTACC | TCTTCCTCTTCATCATCTTCATCA |
| BIM | CATCGCGGTATTCGGTTC | GCTTTGCCATTTGGTCTTTTT |
| PUMA | GACCTCAACGCACAGTACGA | GAGATTGTACAGGACCCTCCA |
| MAX | CCAGCAAGATATTGACGACCT | TTCTCCAGTGCACGGACTT |
| SUPT5H | GTATGAGGACGAGGACCAGTG | GAACGATCTTCATCCAGGACA |
| KAT2A | GTGCTGTCACCTCGAATGAG | TGGAGAAACCCTGCTTTTTGA |
| KAT5 | CGTAAGAACAAGAGTTATTCCCAG | GTCTTCCGTTGATTCTTTCTCC |
| RUVBL1 | CTGTGTCATCAGAGGCACTGA | AAGTTCACTGATCTCTTCGACATG |
| RUVBL2 | CGAGGAAGAAGATGTGGAGA | CACTTCTGTACCCTTGCGTT |
| TRRAP | CAGGAAGTGAAACGCTTTAG | GTCTTCAGAAGGTTCACACACC |
| VDAC3 | TAATTTCGCCCTGGGTTACA | AAATTCAGTGCCATCGTTCA |
| GAPDH | AGCCACATCGCTCAGACAC | GCCCAATACGACCAAATCC |
| PGK1 | CAGCTGCTGGGTCTGTCAT | GCTGGCTCGGCTTTAACC |
| TBP | CCCATGACTCCCATGACC | TTTACAACCAAGATTCACTGTGG |
